# Extracellular matrix proteolysis maintains synapse plasticity during brain development

**DOI:** 10.1101/2025.02.27.640672

**Authors:** Haruna Nakajo, Ran Cao, Supriya A. Mula, Justin McKetney, Nicholas J. Silva, Muskaan Shah, Indigo V. L. Rose, Martin Kampmann, Danielle L. Swaney, Christoph Kirst, Anna V. Molofsky

## Abstract

Maintaining a dynamic neuronal synapse pool is critical to brain development. The extracellular matrix (ECM) regulates synaptic plasticity via mechanisms that are still being defined and are studied predominantly in adulthood. Using live imaging of excitatory synapses in zebrafish hindbrain we observed a bimodal distribution of short-lived (dynamic) and longer-lived (stable) synapses. Disruption of ECM via digestion or brevican deletion destabilized dynamic but not stable synapses and led to decreased synapse density. Conversely, loss of matrix metalloproteinase 14 (MMP14) led to accumulation of brevican and increased the stable synapse pool, resulting in increased synapse density. Microglial MMP14 was essential to these effects in both fish and human iPSC-derived cultures. Both MMP14 and brevican were required for experience-dependent synapse plasticity in a motor learning assay. These data, complemented by mathematical modeling, define an essential role of ECM remodeling in maintaining a dynamic subset of synapses during brain development.

## Introduction

Neuronal synapse numbers increase markedly during brain development and undergo a prolonged period of experience dependent refinement that shapes adult brain function^1^. In humans, prefrontal cortical synapses increase throughout early childhood and are subsequently pruned in adolescence^2,3^ and alterations in synapse plasticity are implicated in neurodevelopmental diseases^4,5^. The extracellular matrix (ECM) is a lattice of sugars and glycoproteins that fills the extracellular spaces of the brain, which comprise up to 20% of brain volume^6^. The ECM is also a critical regulator of synaptic plasticity^7,8^. Much of the evidence for this view comes from studies that enzymatically digest the ECM in adulthood. These have found that ECM digestion can reopen plasticity in cortical circuits^9–11^, impair learning and memory^12,13^, and promote recovery from central nervous system (CNS) injury^14,15^. However, ECM protein composition and glycosylation patterns change over the course of development, and some of these changes may be linked to the closure of critical periods^16,17^. Taken together, this suggests that the ECM has a unique molecular composition in development compared to adulthood, and could have temporally distinct roles.

The abundance and composition of ECM depends both on its structural components as well as enzymes that digest and remodel it. Many structural ECM proteins impact CNS synaptic function, including chondroitin sulfate proteoglycans (CSPGs) such as aggrecan and brevican^9,18–20^, link proteins^21–23^, and tenascins^24,25^, which collectively assemble onto the hyaluronan backbone in the brain’s extracellular space. However, the endogenous mechanisms that remodel the ECM are less understood, in part because many studies have bypassed those mechanisms by non-physiologic injections of enzymes. ECM remodeling enzymes include matrix metalloproteinases of the MMP, ADAM, and ADAMTS families^26–28^. Among these is the matrix metalloproteinase MMP14, which is amongst a small number of MMPs that are membrane bound, potentially enabling ECM remodeling in a cell contact dependent manner.

Microglia, the innate immune cells of the CNS parenchyma, promote ECM remodeling in response to immune cues such as Interleukin-33 and CX3CL1^29–34^. Live imaging in adult mice or slice preparations from developing brains shows that microglial contact induces spine formation and spine plasticity^35,36^, a phenotype that mirrors the effect of localized injection of ECM degrading enzymes^9^. However, studies of synapse, ECM, and microglial dynamics in the intact developing brain have been limited by imaging constraints in developing mammals. In contrast, zebrafish (*Danio rerio*) are a vertebrate model system that develops ex utero and is transparent up to 14 days post fertilization (dpf), enabling dynamic imaging of core neurodevelopmental processes including synapse formation^37–39^. We recently characterized a population of microglia in the zebrafish hindbrain that interact with synapses and recapitulate core morphological and molecular features of mammalian synapse-associated microglia^40^, facilitating studies of microglia-synapse interactions during development. Although ECM is generally thought to restrict plasticity in the adult brain, it is less clear how the very different ECM milieu of the developing brain impacts synapse plasticity and circuit function, and whether these processes are affected by microglia.

Here, we show that the structural ECM protein brevican and the matrix metalloproteinase MMP14 are key regulators of developmental synapse plasticity. Via time-lapse live imaging in the developing zebrafish hindbrain, we identify two pools of excitatory synapses with distinct kinetics. One pool comprises the majority of synapses and consists of long-lived synapses (days). However, a substantial minority of synapses are quite dynamic, often lasting only a few hours. We show that loss of brevican or ECM digestion reduces excitatory synapses by preferentially destabilizing this short-lived pool of synapses. We also show that MMP14, which is expressed by microglia, promotes remodeling of ECM proteins including brevican in both zebrafish and human induced pluripotent stem cell (iPSC) derived cultures, and that loss of MMP14 increases the proportion of synapses in a stable state. Finally, we show that these pathways are required for experience-dependent synaptic plasticity in a forced swim model of motor learning. Together, these data suggest that a key function of ECM remodeling during development is to maintain a dynamic pool of synapses to enable experience-dependent adaptation.

## Results

### Characterization of synapse dynamics and extracellular matrix (ECM) in the developing brain

The brain’s perisynaptic ECM consists of a protein scaffold of chondroitin sulfate proteoglycans (CSPGs). Brevican is an abundant and brain-specific CSPG that plays important roles in synapse development and plasticity^18,41^. CSPGs are connected by linker proteins such as tenascins and assembled onto a backbone of hyaluronan sugar chains (**Fig. 1a**). We quantified brevican and hyaluronan between 7-90 days post fertilization (dpf), spanning the equivalent of mammalian embryonic development through adulthood. We focused on the hindbrain region because it is synapse rich and accessible to live imaging, and contains a population of synapse-associated microglia which we have previously characterized^40^. Brevican was quantified by immunostaining and hyaluronan was visualized using the reporter fish *Tg(ubi:ssncan-GFP)* which expresses the hyaluronan binding domain of neurocan fused to GFP^42^ (**Fig. 1b, Extended Data Fig. 1a**). We found that hyaluronan expression was mostly stable over development, whereas brevican increased progressively until 60 dpf and was 3-fold enriched in the adult vs. developing brain (**Fig. 1c**), consistent with studies in rodents^43,44^. In mammals, dense-ECM structure around neurons called perineuronal nets (PNN) have critical roles in regulating synapse plasticity and neural circuit functions in adult brain^45,46^. We found that adult zebrafish have net-like brevican structures that were not observed in development (**Extended Data Fig. 1b**). These results suggest a distinct composition and structure of the developing brain ECM compared to adulthood.

**Figure 1:**
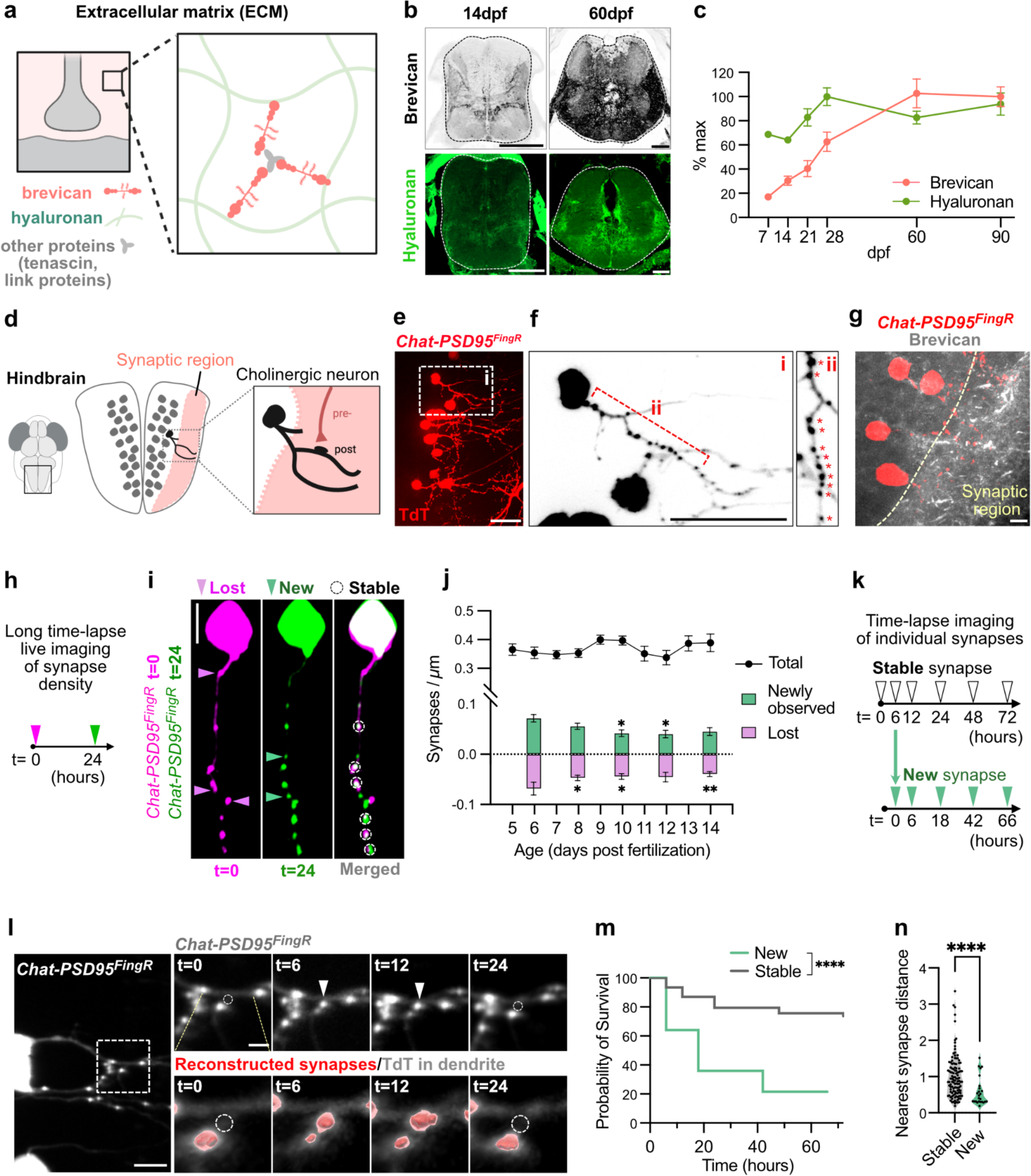
Characterization of synapse dynamics and extracellular matrix (ECM) in the developing brain. a) Schematic of the perisynaptic ECM which consists of a hyaluronan sugar matrix connected by proteoglycans including brevican, linking proteins like tenascins, and others. b) Representative images of zebrafish hindbrain at 14 and 60 days post fertilization (dpf) showing brevican (antibody staining) and hyaluronan labeled with a genetically encoded sensor *ubi:ssncan-GFP*. Scales: 100 µm. c) Quantification of fluorescence intensity of brevican and hyaluronan (*ubi:ssncan-GFP*) over development from 7-90 dpf. Mean fluorescence intensity for brevican and hyaluronan were normalized to the maximum intensity measured for each (90 dpf and 28 dpf, respectively). Brevican quantification, number of fish: 7 dpf, n=9; 14 dpf, n=8; 21 dpf, n=8; 28 dpf, n=7; 60 dpf, n=7; 90 dpf, n=6. For hyaluronan, number of fish: 7 dpf, n=11; 14 dpf, n=10; 21 dpf, n=6; 28 dpf, n=10; 60 dpf, n=7; 90 dpf, n=3. d) Schematic of zebrafish larval hindbrain. Gray dots indicate cell bodies and the synaptic region is shown by pink. A cholinergic neuron with a cell body and dendrites is shown by black. e) Dorsal view of zebrafish hindbrain at 10 dpf shows sparsely labeled cholinergic neurons expressing a TdTomato (TdT) tagged FingR construct that binds to the excitatory postsynaptic marker PSD-95. *Tg(chata:gal4);Tg(zcUAS:PSD95.FingR-TdT-CCR5TC-KRAB(A)),* hereafter abbreviated *Chat-PSD95^FingR^*. Scale: 20 µm. f) Strategy for quantification of synapses using *Chat-PSD95^FingR^*. Insets of region in e shows (i) several sparsely labeled neurons and (ii) a dendritic segment from one cholinergic neuron with synapses indicated by asterisks. Scales: 20 µm. g) Immunostaining for brevican protein and *Chat-PSD95^FingR^*at 14 dpf. Synaptic region is indicated in the image. Scale: 5 µm. h) Schematic of 24 hours time lapse imaging assay to quantify changes of synapse density. i) Representative images show a single *Chat-PSD95^FingR^*dendrite imaged at 7 dpf (t=0) and 8 dpf (t=24). Pink arrowheads: synapses present at t=0 and absent at t=24 “lost synapses”. Green arrowheads: synapses that appear at t=24 “new synapses”. White circles: present at both time points. Scale: 5 µm. j) Quantification of total excitatory synapse density and dynamics over the live imaging window of hindbrain development (5-14 dpf), based on *Chat-PSD95^FingR-TdT^* puncta normalized to µm of dendrite length. Black line indicates static synapse density per day (p=0.3437, One-way ANOVA). Pink bars indicate lost synapses (F_(4,78)_ p=0.0040**, One-way ANOVA). Green bars indicate new synapses (F_(4,78)_ p=0.018*, One-way ANOVA). Asterisks in the figure represent results of Tukey’s multiple comparisons with respect to the 5-6 dpf. Number of fish: 5-6 dpf, n=15; 7-8 dpf, n=18; 9-10 dpf, n=18; 11-12 dpf, n=15; 13-14 dpf, n=17. Results from individual fish in Extended Data Fig. 2b. k) Schematic of time lapse imaging to determine the fate of individual synapses at the indicated timepoints. Synapses at t=0 were defined as “stable” synapses. Synapses born between t=0 and t=6 were defined as “new” synapses and subsequently followed with the 6 hour timepoint set as t=0 (lower timecourse). Experiments were performed at 10-12 dpf. l) Representative image of a single excitatory synapse imaged at t=0, 6, 12, and 24 shows a “new synapse” born between t=0 and t=6, which disappeared between t=12 and t=24. Left shows a low power image of a *Chat-PSD95^FingR^* cholinergic neuron. Dashed square indicates inset. Inset shows raw fluorescence (top) and fluorescence overlaid with 3D reconstruction of synapses (bottom). Arrowheads: newborn synapse. Circle: site of newborn synapse. Scale: 5 µm in low power image and 2 µm in inset. m) Kaplan-Meier plot of survival of individual synapses over time (Data from n=19 fish, n=427 stable synapses and n=25 new synapses). n) Quantification of the distance from the nearest synapse for stable and new synapses at t=6 (Stable, n=116 inter-synapse intervals from n=14 fish; New, n=25 inter-synapse intervals from n=14 fish; p<0.0001****, Welch’s t-test, performed for synapses). The inter-synapse intervals were normalized by the mean of stable synapses for each cell. Values were plotted as mean ±SEM. ****: p<0.0001; **: p<0.01; *: p<0.05; ns: not significant.

We next established tools to examine synapses in developing hindbrain, focusing on cholinergic neurons which can be genetically targeted with the promoter of the choline acetyltransferase gene *chata*^47^. The larval hindbrain has a distinct cell-body rich region where most neuronal soma are located, including hindbrain cholinergic neurons. The dendrites of these neurons are in a synapse rich neuropil where they receive pre-synaptic input from other neurons (**Fig. 1d**). To visualize excitatory synapses onto cholinergic neurons, we used a transgenic system based on FingR technology (fibronectin intrabodies generated by mRNA display), which can report on endogenous synaptic proteins with minimal impact on neural function, protein trafficking, or expression^48,49^. We established a transgenic (Tg) line expressing FingR targeted to post-synaptic density protein 95 (PSD95) using the GAL4/UAS system to sparsely label neurons (**Fig. 1e,f**)^47–49^. This transgenic fish *Tg(chata:gal4); Tg(zcUAS:PSD95.FingR-TdT-CCR5TC-KRAB(A))* is hereafter referred to as *Chat-PSD95^FingR^*. This line had sufficient cytoplasmic signal to identify neuronal soma and processes, as well as distinct excitatory synaptic puncta (**Fig. 1f**, asterisks). We observed that brevican diffusely surrounded neurons and their synapses in the developing hindbrain (**Fig. 1g**).

Synapses are dynamic during early development and become progressively more stable in the mature brain^50^. We used *Chat-PSD95^FingR^* fish to quantify both synapse numbers and the dynamics of synapses during development. We quantified excitatory synapse numbers using single images from fixed or live-imaged brains as shown in **Fig. 1f**. To quantify synapse dynamics, we established time-lapse assays to visualize synaptic turnover by imaging the same neuron at two time points separated by 24 hours (t=0 and t=24; **Fig. 1h**). Using these two approaches, we could quantify both synapse density at each imaging day, as well as synapses that formed or were removed during the preceding 24 hour window (**Fig. 1i**).

We first quantified total synapse density and dynamics between 5-14 dpf, which spans the live imaging window. To do this, we performed serial imaging of *Chat-PSD95^FingR^* in groups, whereby one group of larvae was imaged at 5 dpf and again at 6 dpf, a second group was imaged at 7 dpf and again at 8 dpf, and so on. We found that excitatory synapse density was largely stable over this imaging period (**Fig. 1j, black line**). Neuronal dendrite length and branching was also stable (**Extended Data Fig. 2a**). However, despite stable overall synapse density, time lapse imaging showed that 24 ±1 % of synapses were replaced in each 24 hour period between 5-14 dpf (**Fig. 1j**, pink and green bars, and **Extended Data Fig. 2b,c).** We termed these “newly observed” and “lost” synapses. The term “newly observed” was meant to clarify that these numbers reflected an average over a 24 hour window, and might not capture more transient “new” synapses, which were examined in more detail below. While synapse turnover was mildly decreased at later time points (after 9-10 dpf), overall, our data suggest a remarkable amount of synaptic turnover throughout this developmental window.

While the above data suggest that synapses are continuously replaced, our 24 hours imaging window could fail to capture synapses that are transient within that 24 hours period. Studies suggest that while some synapses are stable over the life of the organism^51,52^, others live only a few hours^39^. To investigate synapse lifetime in our system, we tracked individual synapses by serial time lapse imaging over 72 hours. Synapses present at the start of the imaging window (t=0) were defined as “stable”, whereas synapses that appeared between t=0 and t=6 were defined as “new” (**Fig. 2k**). We found that 74% of stable synapses were still present at t=72. In contrast, fewer than 20% of new synapses born between t=0 and t=6 were still present at the end of the imaging window (**Fig. l l,m**). We also found that new synapses tended to form in proximity to other synapses (**Fig. 2n**), reminiscent of findings in adult mouse motor cortex^53^. PSD95 diameter between newborn and stable synapses was not different (**Extended Data Fig. 2d**). These findings suggest that while a majority of synapses are stable (lifetime >72 hours), a smaller proportion of synapses live only a few hours.

**Fig. 2:**
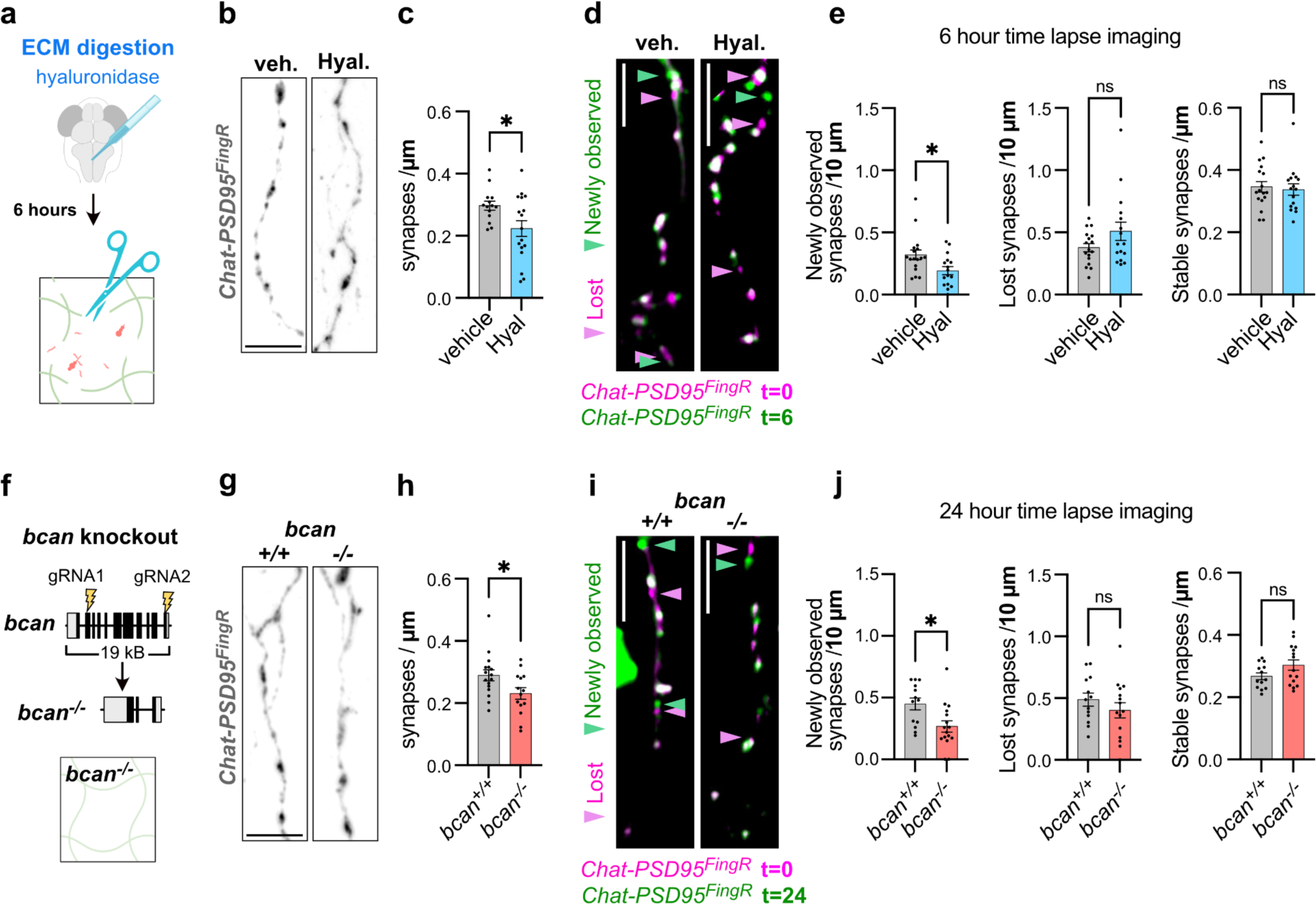
Extracellular matrix depletion preferentially destabilized newborn synapses. a) Schematic of strategy for ECM digestion by hyaluronidase injection into the hindbrain ventricle. Tissues fixed or imaged 6 hours after injection with hyaluronidase or vehicle (PBS). b) Representative images of *Chat-PSD95^FingR^* dendrites at 14 dpf from fixed section stained for ΤdT after vehicle or hyaluronidase injection. Scale: 5 µm. c) Quantification of synapse density at 14 dpf (Chat-PSD95^FingR^ puncta) per µm of dendrite length. Dots represent means per fish, with at least 1 dendrite quantified per fish (vehicle, n=14 fish; hyaluronidase, n=17 fish; p=0.020*, Welch’s t-test). d) Representative merged images of *Chat-PSD95^FingR^*collected before injection (t=0, pink) and 6 hours after (t=6, green) hyaluronidase or vehicle injection. Overlap of pink and green appears white. Pink arrowheads: lost synapses, green arrowheads: newly observed synapses. Experiment performed at 10-12 dpf. Non-merged images in Extended Data Fig. 3h. Scale: 5 µm. e) Quantification of newly observed, lost, or stable synapses between t=0 and t=6 after hyaluronidase vs. vehicle injection (vehicle, n=17 fish; Hyal, n=16 fish; p=0.017* for newly observed, p=0.117 for lost, p=0.722 for stable, Welch’s t-test). f) Schematic of generation of brevican knock-out (*bcan^-/-^*) fish by CRISPR genome editing. Guide RNAs targeting exon 3 and exon 14 were injected to delete 19 kbp of the *bcan* gene. The truncation resulted in loss of *bcan* mRNA and brevican protein (see Extended Data Fig 3e,f). g) Representative images of *Chat-PSD95^FingR^* dendrites at 14 dpf from fixed section stained for ΤdT from *bcan^+/+^*vs. *bcan^-/-^* fish. Scale: 5 µm. h) Quantification of synapse density at 14 dpf (Chat-PSD95^FingR^ puncta) per µm of dendrite length. Dots represent means per fish, with at least 1 dendrite quantified per fish (*bcan^+/+^*, n=16 fish; *bcan^-/-^*, n=14 fish; p=0.031*, Welch’s t-test). i) Representative merged images of *Chat-PSD95^FingR^* collected from *bcan^+/+^* and *bcan^-/-^* at t=0 (pink) and t=24 (green). Pink arrowheads: lost synapses, green arrowheads: new synapses. Experiment performed at 10-12 dpf. Non-merged images in Extended Data Fig. 3i. Scale: 5 µm. j) Quantification of newly observed, lost and stable synapses between t=0 and t=24 *bcan^+/+^* vs. *bcan^-/-^* fish (*bcan^+/+^*, n=13 fish; *bcan^-/-^*, n=15 fish; p=0.010* for newly observed, p=0.288 for lost, p=0.085 for stable, Welch’s t-test). Values were plotted as mean ±SEM. *: p<0.05; ns: not significant.

### Extracellular matrix depletion preferentially destabilized newborn synapses

To define how ECM impacts developing synapses, we examined both static synapse numbers and synapse dynamics using two independent approaches to deplete the ECM. First, we injected hyaluronidase into the hindbrain ventricle of 14 dpf larvae to acutely digest the hyaluronan scaffold (**Fig. 2a**) and performed analysis six hours after hyaluronidase or vehicle (PBS) injections. We chose these timepoints to capture the maximal reduction in ECM density after hyaluronidase, which decreased by six hours post injection (hpi) and had recovered to baseline levels by 24 hpi (**Extended Data Fig. 3a-c**). We found that the acute ECM depletion with hyaluronidase significantly reduced excitatory synapse density at 14 dpf as measured by Chat-PSD95^FingR^ puncta (**Fig. 2b,c**), as well as the presynaptic marker SV2 (**Extended Data Fig. 3d**). We also performed time lapse imaging before and six hours after hyaluronidase or vehicle injections (**Fig. 2d,e, Extended Data Fig. 3e**). ECM digestion affected synapse dynamics: we observed fewer newly observed synapses relative to vehicle control, whereas the number of lost and stable synapses were unchanged (**Fig. 2e**).

To test the impact of the ECM using an independent approach, we generated brevican knockout (*bcan^-/-^*) fish using CRISPR genome editing (**Fig. 2f**). We confirmed complete loss of *bcan* mRNA and brevican protein (**Extended Data Fig. 3f,g**). Similar to our results after acute enzymatic depletion, we found that brevican-deficient larval fish had less Chat-PSD95^FingR^ excitatory synapse density relative to *bcan^+/+^* controls (**Fig. 2g,h**), supported by staining for the presynaptic marker SV2 (**Extended Data Fig. 3h**). This decrease is consistent with a prior study in juvenile mouse hippocampus^18^. Time lapse imaging over 24 hours showed that this overall loss of synapses corresponded with a reduced number of newly observed synapses, but no changes in the lost or stable synapse pools (**Fig. 2i,j, Extended Data Fig. 3i**). Interestingly, ECM depletion had the opposite effect in adult fish (60 dpf), whereby both hyaluronidase injection and brevican knockout increased synapse density (**Extended Data Fig. 4**). These data suggest that the ECM may differentially impact developing vs. adult synapses. In development, loss of ECM preferentially impairs a short-lived synapse pool and leads to an overall loss of synapses.

### The microglial metalloproteinase Mmp14b restricted ECM accumulation and stabilized newborn synapses

We previously showed that microglia promote synapse plasticity in the adult mouse brain by restricting accumulation of ECM^29^. We therefore examined whether microglia interact with synapses in developing hindbrain and whether they impact ECM accumulation. To visualize microglia-synapse interactions we crossed our *Chat-PSD95^FingR^* fish to a reporter line that expresses membrane-targeted GFP in macrophage-lineage cells, including microglia (*Tg(mpeg:GFP-CAAX)* hereafter “*mpeg-GFP*”).

Zebrafish hindbrain has a distinct population of synapse-associated microglia that molecularly and morphologically resemble mammalian microglia^40^. We observed that microglia in the hindbrain clustered near the synaptic region^40^ (**Extended Data Fig. 5a,b**). Using live imaging, we observed frequent contacts between microglia and excitatory synapses over the 1 hour imaging window (10-12 dpf, **Extended Data Fig. 5c,d, Supplemental Video 1**). Out of 53 contacts examined, none of these contacts were associated with subsequent disappearance of the synapse within the time window of imaging. ECM digestion with hyaluronidase modestly increased the frequency of microglia-excitatory synaptic contacts without altering overall process motility (**Extended Data Fig. 5e-g**), suggesting that ECM could restrict microglial contact with synapses in some settings. These data demonstrate microglia contact excitatory synapses in the developing brain.

To investigate if microglia impact ECM density in the developing brain, we conditionally ablated microglia by expressing nitroreductase (NTR) in mpeg+ cells, which causes them to undergo apoptotic cell death in the presence of metronidazole (**Fig. 3a**; (“*Tg(mpeg:gal4);Tg(UAS:NTR-mCherry)”).* This strategy led to a partial (46±12%) reduction in total microglia after 24 hours (**Extended Data Fig. 6a**), likely because not all microglia expressed the NTR-mCherry construct. Fish with partial ablation of microglia had increased brevican signal in the synaptic region (**Fig. 3b,c**). Metronidazole alone did not alter brevican intensity in non-transgenic siblings (**Extended Data Fig. 6b**). These results suggest that microglia restrict ECM accumulation in the developing hindbrain, consistent with their role in the adult brain^29,54^.

**Figure 3:**
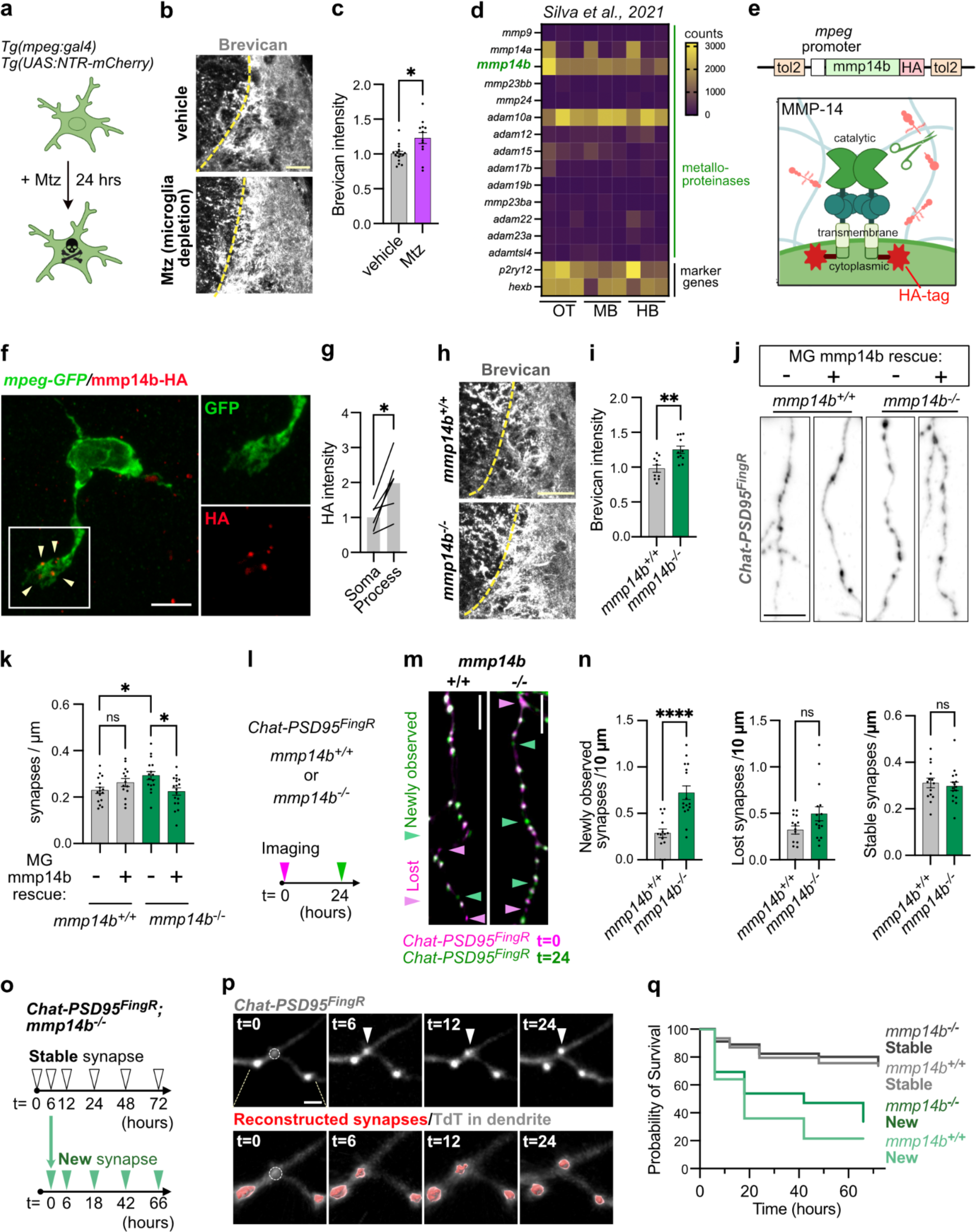
The microglial metalloproteinase Mmp14b restricted ECM accumulation and stabilized newborn synapses. a) Schematic of microglial ablation by adding metronidazole (Mtz) to fish water in fish that express nitroreductase (*ntr*) in microglia and macrophages (*Tg(mpeg:gal4);Tg(UAS:NTR-mCherry)*). b) Representative images of brevican in synaptic region after vehicle (DMSO) or microglial ablation with 5mM metronidazole (Mtz) in fish water for 24 hours. Experiment at 14 dpf. Scale: 20 µm. c) Quantification of brevican intensity in synaptic region normalized to mean of vehicle control (vehicle, n=15 fish; Mtz, n=11 fish; p=0.043*, Welch’s t-test). d) Heatmap of absolute expression (counts) of *mmp*, *adam*, and *adamts* metalloproteinase genes in zebrafish microglia from bulk RNAsequencing at 28 dpf in optic tectum (OT), midbrain (MB) and hindbrain (HB). Marker genes *hexb* and *p2ry12* shown for comparison. Reanalyzed from ^40^. e) Schematic of a dimer of the membrane-associated metalloproteinase MMP14 and design of the *mmp14b-HA* expression construct for epitope tagging of the C-terminal end with HA. Catalytic domain, transmembrane domain and cytoplasmic domain are indicated. f) Mmp14b-HA transgene expression in a *mpeg-GFP* microglia expressing mpeg:mmp14b-HA. Inset shows HA signals colocalized with microglial processes (arrow heads). Scale: 5 µm. g) Quantification of Mmp14b-HA intensity in microglial processes vs soma. Lines connect data from the same microglia (n= 6 microglia from 3 fish; p=0.015*, Paired t-test per cell). h) Representative images of brevican staining in synaptic regions of *mmp14b^-/-^* and *mmp14b^+/+^* control at 14 dpf. Scale: 20 µm. i) Quantification of brevican intensity in synaptic region normalized to mean of *mmp14b^+/+^* control (*mmp14b^+/+^*, n=10 fish; *mmp14b^-/-^*, n=11 fish; p=0.0014**, Welch’s t-test). j) Representative images of microglia-specific Mmp14b rescue experiment. *Chat-PSD95^FingR^ ;mmp14b^+/+^* and *;mmp14^-/-^* fish were injected with mpeg:mmp14b-HA construct at one-cell stage embryos and analyzed at 14 dpf. Scale: 5 µm. k) Quantification of synapse density (Chat-PSD95^FingR^ puncta per µm of dendrite length) in *mmp14b^+/+^* and *mmp14^-/-^* with or without microglial-specific rescue with mpeg:mmp14b-HA. Dots show means per fish from at least 1 dendritic segment analyzed per fish (*mmp14^+/+^* no rescue, n=16 fish; *mmp14b^+/+^*rescue, n=14 fish; *mmp14b^-/-^* no rescue, n=16 fish; *mmp14b^-/-^*rescue, n=18 fish; Two-way ANOVA, F_(1,60)_ interaction effect p=0.0022**, asterisks show Tukey’s multiple comparisons). l) Schematic of time lapse imaging to measure synapse turnover in *Chat-PSD95^FingR^ ;mmp14b^+/+^* and *;mmp14b^-/-^* fish. m) Representative merged images of synapses in *Chat-PSD95^FingR^;mmp14b^-/-^* and ;*mmp14b^+/+^*fish at t=0 (pink) and t=24 (green). Experiments at 10-12 dpf. Pink arrowheads: lost synapses, green arrowheads: newly observed synapses. Non-merged images in Extended Data Fig. 8d. Scale: 5 µm. n) Quantification of newly observed, lost, and stable synapses between t=0 and t=24 in *mmp14b^+/+^* vs *mmp14b^-/-^* fish (*mmp14b^+/+^*, n=14 fish; *mmp14b^-/-^*, n=16 fish; p<0.0001**** for newly observed, p=0.058 for lost, 0.61 for stable, Welch’s t-test). o) Schematic of time lapse imaging to determine the fate of individual synapses in *mmp14b^-/-^* fish. Synapses at t=0 were defined as “stable” synapses. Synapses born between t=0 and t=6 were defined as “new” synapses and subsequently followed with the 6 hour timepoint set as t=0 (lower timecourse). Experiments performed at 10-12 dpf. p) Representative image of a single excitatory synapse imaged at t=0, 6, 12, and 24 shows a “new synapse” born between t=0 and t=6 in *mmp14b^-/-^* fish. Raw fluorescence images on top and fluorescence overlaid with 3D reconstruction of synapse on bottom. Arrowheads: newborn synapse. Circle: site of newborn synapse. Scale: 2 µm. q) Kaplan-Meier plot of survival of individual synapses over time in *mmp14b^+/+^* control (from Fig. 1m) and *mmp14b^-/-^* fish (For *mmp14b^-/-^*, Data from n=15 fish, n=331 stable synapses and n=26 new synapses). Values were plotted as mean ±SEM. ****: p<0.0001; **: <0.01; *: p<0.05; ns: not significant.

To determine potential microglial regulators of the ECM, we screened zebrafish microglia for expression of the major classes of metalloproteinases, which include genes in the MMP (matrix metalloproteinase), ADAM (a disintegrin and metalloproteinase) and ADAMTS (a disintegrin and metalloproteinase with thrombospondin motifs) families^26–28^. We examined expression of all genes in these families from our previously published bulk RNA sequencing of purified zebrafish microglia at 28 dpf^40^, which included samples from optic tectum (OT), midbrain (MB), and hindbrain (HB). We identified 14 proteinases with detectable expression (>100 counts in >1 replicate). Among these, the top expressed metalloproteinases were *mmp14b* and *adam10a*, both of which were detected at levels comparable to canonical microglial genes like *p2ry12* and *hexb* (**Fig. 3d**). Interestingly, both *mmp14b* and *adam10a* are membrane bound, potentially enabling them to cleave proteins in a cell contact dependent manner^55,56^. We previously identified mouse *Mmp14* as a top candidate in a screen for ECM remodeling enzymes expressed in mouse microglia^29^. We confirmed *mmp14b* expression in 60±8 % of hindbrain microglia at 14 dpf by RNA *in situ* hybridization (**Extended data Fig. 7a-c**). *Mmp14b* was also detected in astrocytes but was rarely detected in neurons (**Extended Data Fig. 7d,e).** The probe was specific to *mmp14b*, as no hybridization was observed in *mmp14b* knockout fish (*mmp14b^-/-^*^57^; **Extended Data Fig. 7f).**

MMP14 can directly cleave ECM proteins and activate other metalloproteinases through cleavage of their pro-domains^55^. To directly observe the subcellular localization of MMP14 in microglia, we generated a construct that expresses Mmp14b with an HA epitope tag at its C-terminal end driven by the *mpeg* promoter (*mpeg:mmp14b-HA*; **Fig. 3e**). We injected this transgenic construct into one-cell stage embryos and examined expression at 14 dpf. We observed Mmp14b-HA protein preferentially localized at microglial processes (**Fig. 3f,g**). Thus, microglial MMP14 could potentially influence microglial process function, including their motility, contact with synapses, or cleavage of ECM in a contact-dependent manner.

To determine the function of Mmp14b *in vivo* we examined *mmp14b^-/-^*fish which harbor a deletion from exon 6 to 9 of the *mmp14b* gene^57^. These fish have complete loss of *mmp14b* transcript (**Extended Data Fig. 7f**) without a compensatory upregulation of the paralog gene *mmp14a*^57^, which we detected only sporadically in zebrafish microglia (**Fig. 3d**). While MMP14 deficient mice show visible dysmorphism around 5 days after birth^58^, fish lacking *mmp14b* are viable and fertile with only a small population of adult fish showing skeletal defects^57^. *mmp14b^-/-^* fish had normal numbers of microglia (**Extended Data Fig. 8a**), however, their microglia had shorter processes, as measured by total process length (**Extended Data Fig. 8b**). *mmp14b^-/-^* fish also had increased brevican in the synaptic region (14 dpf; **Fig. 3h,i**), indicating that Mmp14b restricts ECM accumulation in the developing hindbrain.

To quantify the effect of Mmp14b on excitatory synapses, we compared *mmp14b^-/-^;Chat-PSD95^FingR^* to control *mmp14b^+/+^;Chat-PSD95^FingR^*fish. We found an increase in Chat-PSD95^FingR^ excitatory synapse density in the *mmp14b^-/-^* fish compared to controls (14 dpf; **Extended Data Fig. 8c**) and an increase in the presynaptic marker SV2 (**Extended Data Fig. 8d**), which was the opposite of what we had observed after ECM depletion (**Figure 2**). Since *Mmp14b* was also detected in astrocytes, we performed a microglia specific rescue of *mmp14b* to determine whether this was sufficient to restore synaptic density. We injected the *mpeg:mmp14b-HA* construct from Fig. 3e-f into *mmp14b^-/-^;Chat-PSD95^FingR^* fish and *mmp14b^+/+^;Chat-PSD95^FingR^* fish, and quantified excitatory synapses at 14 dpf. We found that microglial expression of Mmp14b was sufficient to rescue excitatory synapse density to control levels in *mmp14b^-/-^* fish, but did not increase excitatory synapse density in *mmp14b^+/+^*, suggesting that microglial mmp14b is sufficient to regulate synapse numbers, and furthermore, that our rescue was not a gain-of function effect (**Fig. 3j,k**). Taken together, these results suggested that microglial Mmp14b restricts accumulation of ECM and excitatory synapses.

To examine the effect of *mmp14b* on synapse dynamics, we performed time lapse imaging in *mmp14b^-/-^;Chat-PSD95^FingR^* versus *mmp14b^+/+^;Chat-PSD95^FingR^* controls (**Fig. 3l**). We found that *mmp14b^-/-^* fish had more newly observed synapses relative to controls, with no change in the number of lost synapses or stable synapses (**Fig. 3m,n, Extended Data Fig. 8e**). These results were the inverse of what we observed in hyaluronidase injected and brevican deficient fish (**Fig. 2**). We also examined the lifetime of new and stable synapses in the *mmp14b^-/-^* fish by time lapse imaging over 72 hours (**Fig. 3o**). We found that in *mmp14b^-/-^* fish, 54% of newborn synapses were still present at 24 hours (vs. 36% in controls) and 33% at 72 hours (vs. 20% in controls; **Fig. 3p,q**; control data from **Fig. 1m**). The survival rate of stable synapses were similar in *mmp14b^+/+^* and *mmp14b^-/-^* (74% vs. 76% at t=72) (**Fig. 3q**). This result suggested that ECM accumulation in the *mmp14b^-/-^* fish could increase the stability of newborn synapses. Taken together, these results suggested that microglial Mmp14b restricts ECM in the developing brain and preferentially destabilizes newly formed synapses relative to stable synapses, thereby restricting total synapse numbers.

### Microglial MMP14 promoted brevican digestion in human cells

Our results in zebrafish indicated opposing effects of microglial MMP14 and brevican on synapse numbers and dynamics. However, they did not address whether these two proteins are functionally linked, and whether these results were applicable to mammals. To test this and further explore the relevance of this pathway across species we established a human tri-culture system of neurons, astrocytes and microglia. *MMP14* is one of the most highly expressed genes in human fetal microglia, comparable in rank to housekeeping genes *AIF1*, *P2RY12,* and *HEXB*^59^. Reanalysis of published datasets revealed strong correlation of gene expression between acutely isolated human fetal microglia and *in vitro* microglia generated from induced pluripotent stem cells (iPSCs; **Fig. 4a**, reanalyzed from^59^). Human microglia express multiple metalloproteinases, of which many are membrane bound, and *MMP14* and *ADAM10* are among the most highly expressed, similar to zebrafish. This suggests that some aspects of microglial MMP14 function can be studied in human cells *in vitro*. To determine the impact of microglial MMP14 on the brain ECM, we co-cultured iPSC-derived neurons^60^ and astrocytes^61^, as these are two of the dominant cell types that produce ECM in the brain. After neuron-astrocyte co-culture and differentiation for 14 days, we added iPSC-derived microglia^62^ and cultured all three cell types together for 24 hours (**Fig. 4b**).

**Fig. 4:**
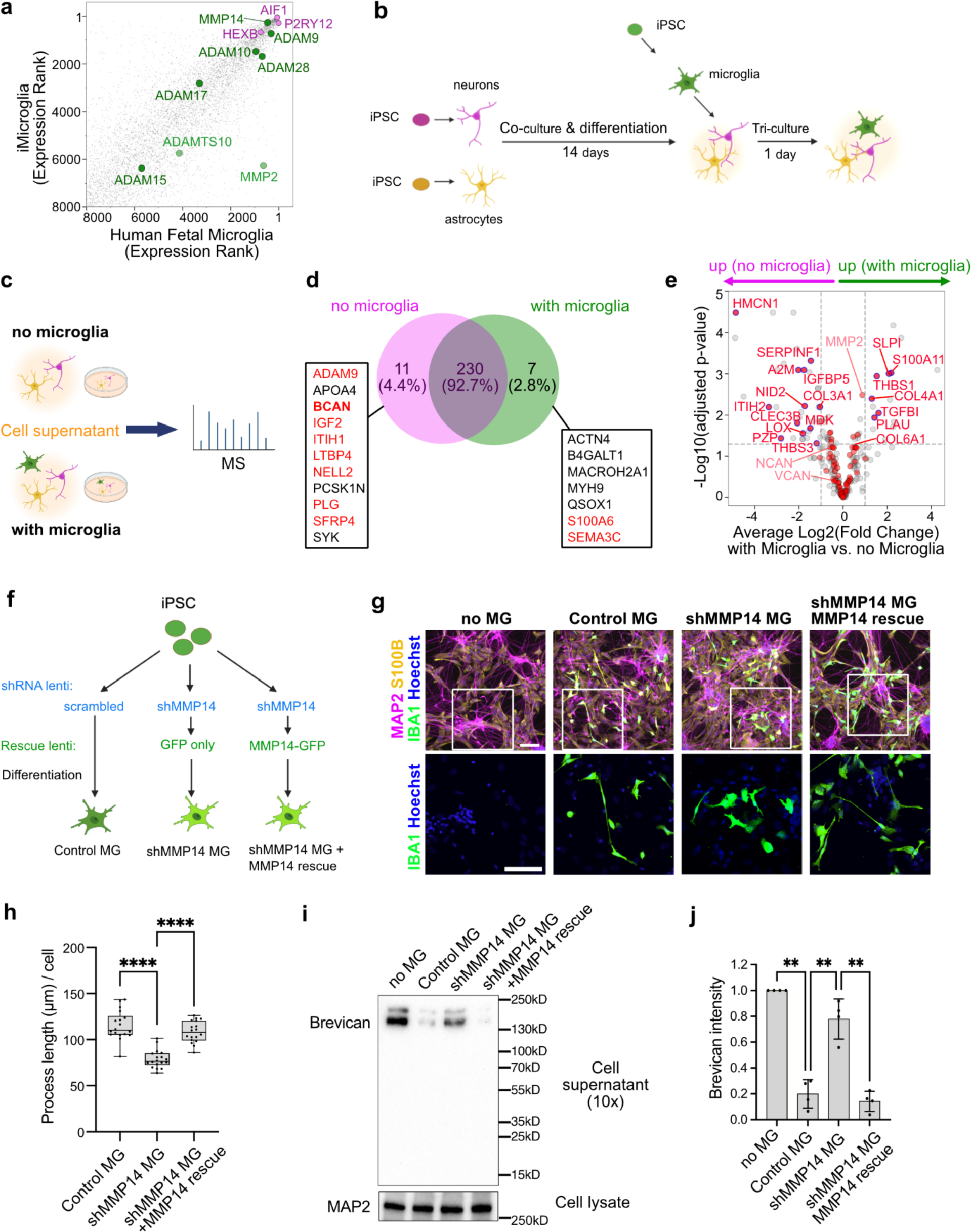
Microglial MMP14 promoted brevican digestion in human cells. a) Scatter plot of gene expression (sorted by rank) of iPSC derived microglia and human fetal microglia. MMPs, ADAMs, ADAMTSs families that have transmembrane domain in dark green; MMPs, ADAMs, ADAMTSs without transmembrane domains in light green; microglial marker genes in violet. Data reanalyzed from the previous study^59^. b) Schematic of the iPSC derived astrocyte-neuron-microglia tri-culture system. c) Schematic of the proteomics analysis with iPSC derived cells. Cell supernatants were collected from co-culture (neurons and astrocytes; no microglia) and tri-culture (neurons, astrocytes and microglia; with microglia) and analyzed. d) Venn diagram of proteins detected in cell culture supernatants with or without microglia. Proteins only detected in the absence of microglia are listed on the left and proteins only detected in the presence of microglia are listed on the right. ECM related proteins as defined by the matrisome database MatrisomeDB2.0^83^ are labeled in red. e) Volcano plot of proteins in the intersection of venn diagram in d. The fold change is calculated by comparing the “with microglia” condition to the “no microglia” condition. Thresholds: p-value<0.05 (horizontal dashed line), and the vertical grey dash lines show where the average log_2_ fold change >1 (vertical dashed lines). ECM related proteins as defined by the matrisome database Matrisome DB2.0^83^ in red. Significantly matrisome proteins are outlined in blue. f) Strategy for MMP14 knockdown (KD) by shRNA (short hairpin RNA interference) with a scrambled shRNA control, and MMP14 rescue by expression of MMP14 fused to GFP, vs a GFP only control in microglia (MG). The rescue construct has silent mutations in the MMP-GFP to prevent knockdown by shMMP14. All constructs were delivered by lentivirus at the iPSC stage and used the ubiquitous promoter EF1a. g) Representative images of the tri-culture system with MMP14 KD and rescue in microglia. Iba1 (microglia), S100β (astrocyte), and neurons (MAP2) stainings are shown. Scale: 100 µm. h) Quantification of the process length of microglia in the tri-culture system with MMP14 KD and rescue. Dots show mean microglial process length per field of view from 18 fields of view over 3 independent experiments. Box-and-whiskers plot: box shows 25-75th percentile, whiskers show min to max. Statistics performed on means per field of view. (Kruskal-Wallis test, asterisks in the figure show results of Dunn’s multiple comparisons). i) Representative image of western blotting for brevican in cell supernatant and MAP2 loading control in cell lysate without microglia or with microglia after MMP14 KD and rescue. j) Quantification of brevican in the cell supernatant without microglia or with microglia after MMP14 KD and rescue. Each dot represents mean brevican intensity normalized to no microglia control from n=4 independent experiments (One-way ANOVA, asterisks in the figure show results of Tukey’s multiple comparisons). Values were plotted as mean ±SEM. ****: p<0.0001; *: p<0.05; ns: not significant.

To assess the ECM production of this co-culture system and the impact of microglia, we performed label-free mass spectrometry on cell culture supernatants at 14 days *in vitro*, comparing neuron-astrocyte co-cultures alone to parallel cultures in which microglia had been added for 24 hours (**Fig. 4c**). We observed robust enrichment for ECM and extracellular proteins in the culture supernatant from both conditions, including the CSPGs brevican, neurocan, and versican (**Fig. 4d,e**). We also observed a substantial change in the proteome after addition of microglia, including loss of brevican detection after addition of microglia (**Fig. 4d**). Interestingly, other ECM proteins were not altered (neurocan, versican), or in some cases increased (COL4A1), suggesting that microglia remodel the ECM in molecularly precise ways. We also noted that MMP2 was detected after addition of microglia. MMP2 is a secreted metalloproteinase that is known to cleave brevican^63^ and can be proteolytically activated by MMP14^55^. These data demonstrate that microglia alter the extracellular proteome and can reduce brevican levels.

To determine the impact of microglial MMP14 on the brain ECM, we repeated tri-culture experiments and knocked down MMP14 by delivering lentiviruses to express short hairpin RNAs (shRNAs) in iPSCs prior to differentiation into microglia (**Fig. 4f**). We identified two shRNA constructs that similarly reduced *MMP14* levels by 50% relative to a scrambled control (**Extended data Fig. 9a**). We subsequently focused on one of these constructs to knock down MMP14. In parallel, we designed a lentiviral construct to express a fusion protein of MMP14 and GFP expressed by the EF1a promoter (**Fig. 4f**). This construct included synonymous mutations in the *MMP14* shRNA binding region to prevent its repression, and was able to restore *MMP14* to control levels at both the mRNA and protein levels (**Extended data Fig. 9b,c**). The rescue construct was compared to a GFP-only expression control, and both these constructs were also delivered to iPSCs prior to differentiation into iMicroglia. These three microglia types (control, MMP14 knockdown, and MMP14 knockdown + rescue) were delivered to our triculture system (**Figure 4f,g**).

Equal numbers of microglia from control, MMP14 knockdown, and MMP14 knockdown + rescue conditions were plated onto neuron-astrocyte co-cultures and microglia were examined 24 hours later. We observed substantial changes in microglial morphology: knockdown of MMP14 led to smaller, rounder microglia with shorter processes, whereas MMP14 rescue increased process length to control levels (**Fig. 4g,h**). Knockdown and rescue of MMP14 did not significantly impact microglial numbers in the tri-culture (**Extended Data Fig. 9d**). These results are similar to the results from *mmp14b^-/-^* fish (**Extended Data Fig. 8a,b**). We then quantified brevican protein levels by western blot. We found that the addition of microglia potently reduced brevican levels relative to the no-microglia control, consistent with our mass spec data. In contrast, adding microglia in which MMP14 was knocked down partly restored brevican levels, and this effect was reversed by MMP14 rescue (**Fig. 4i,j**). WT microglia that did not express any viral constructs showed a similar brevican decrease (**Extended Data fig. 9e**). These data indicate that microglia promote digestion of brevican via MMP14, and that MMP14 impacts microglial morphology. Taken together, these data suggest that MMP14 can promote microglial function in human cells as well as in zebrafish, and that microglial MMP14 either directly or indirectly promotes cleavage and digestion of brevican.

### Brevican and MMP14 regulated neuromotor behaviors

Given these impacts ECM genes on synapse plasticity in hindbrain circuits which include cholinergic motor neurons, we next examined if brevican or Mmp14b impacted sensorimotor behavior. Zebrafish larvae show a startle response upon sudden or potentially threatening visual stimulus, but these responses can diminish with habituation^64,65^, indicating neural circuit adaptation^66^. To study motor activity and habituation we performed a light startle paradigm. Fish were habituated in the dark for 30 min before recording, and spontaneous motor activity was recorded for 10 min both before and after a 40 second train of light pulses (2 sec light/2 seconds dark x10). This paradigm was repeated five times (total time for all five trials: 103 minutes; **Fig. 5a**). Baseline locomotor activity was measured before the first stimulation period. We found that *bcan^-/-^* fish had impaired baseline motility, with less total distance moved and a reduced maximal velocity (**Fig. 5b,c**); however, these baseline parameters were not altered in *mmp14b^-/-^* fish (**Fig. 5d,e**).

**Fig. 5.**
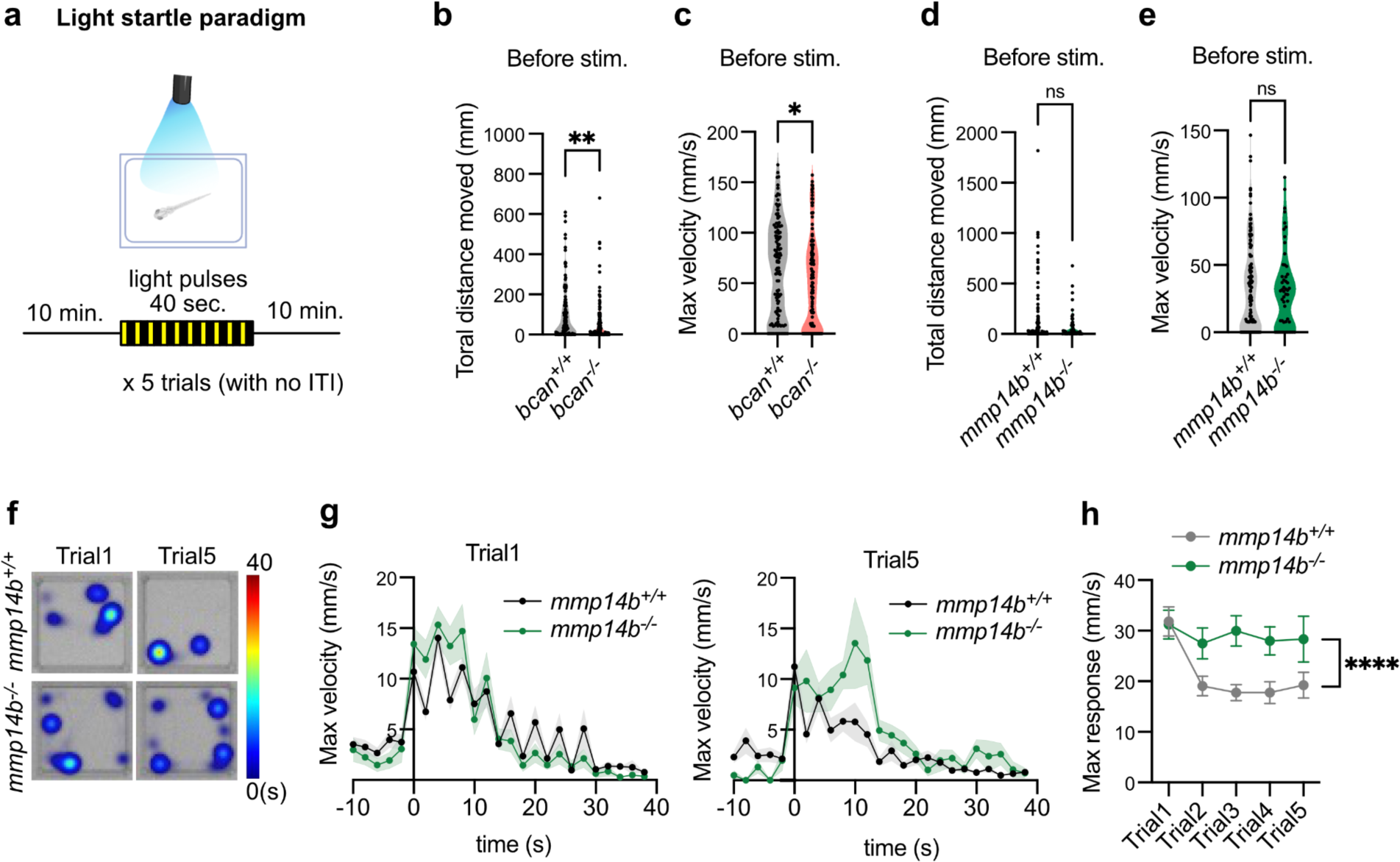
Brevican and MMP14 regulated neuromotor behaviors. a) Schematic of Light startle paradigm performed in this study. For the behavior assay, 14 dpf pigmented fish were used. ITI: inter trial interval. b) Quantification of total distance moved before light stimulus of Trial 1 in the *bcan^-/-^* vs. *bcan^+/+^* control (*bcan^+/+^*, n=120 fish; *bcan^-/-^*, n=128 fish; p=0.011**, Welch’s t-test). c) Quantification of max velocity before light stimulus of Trial 1 in the *bcan^-/-^* vs. *bcan^+/+^*control (*bcan^+/+^*, n=120 fish; *bcan^-/-^*, n=128 fish; p=0.013*, Welch’s t-test). d) Quantification of total distance moved before light stimulus of Trial 1 in the *mmp14b^-/-^* vs. *mmp14b^+/+^* control (*mmp14b^+/+^*, n=115 fish; *mmp14b^-/-^*, n=50 fish; p=0.072, Welch’s t-test). e) Quantification of max velocity before light stimulus of Trial 1 in the *mmp14b^-/-^* vs. *mmp14b^+/+^* control (*mmp14b^+/+^*, n=115 fish; *mmp14b^-/-^*, n=50 fish; p=0.596, Welch’s t-test). f) Representative heatmaps of Trial 1 and Trial 5 from *mmp14^+/+^* and *mmp14^-/-^* fish. The heatmap shows the time spent in each area during the light stimulation period. g) Max velocity for each 2 sec time bin was plotted from 10 sec before the first light stimulus to the end of the last 2 sec dark period. The results from the Trial 1 and Trial 5 are shown (*mmp14b^+/+^*, n=115 fish from trial 1-3, n=75 fish from trial 4-5; *mmp14b^-/-^*, n=50 fish from trial 1-3, n=27 fish from trial 4-5). h) Quantification of maximum response upon light stimuli for each trial (Two-way ANOVA, p-value for genotype). Values were plotted as mean ±SEM.****: p<0.0001; **: p<0.01; *: p<0.05; ns: not significant.

We next examined startle responses in *mmp14b^-/-^* fish compared to *mmp14b^+/+^* controls, and did not test *bcan^-/-^*fish further given their motility impairments. We observed that initial startle in response to light stimuli in trial 1 was not different in *mmp14b^-/-^* fish. However, while control fish habituated to the light by the second trial with a reduction in maximal response speed, *mmp14b^-/-^*showed no habituation throughout the experiment (**Fig. 5f-h**). These data implicate brain ECM as a regulator of neuromotor behavior, as brevican, which is a brain-specific CSPG, was required for normal motility. They also suggest that Mmp14b-dependent ECM remodeling promotes habituation and neural circuit adaptation to stimuli.

### Extracellular matrix accumulation blunted experience-dependent synaptic change

The ability of synapses to remodel in response to experience is a central feature of synaptic plasticity. Given the sensorimotor adaptation deficits in Mmp14b deficient fish, we sought to further probe experience-dependent remodeling with more robust stressors. Involuntary exercise caused by a strong water current (forced swim) is a commonly used assay in fish that can induce a variety of physiologic changes including stress responses and neural plasticity^67,68^, and is analogous to motor learning assays performed in mice^52^. To define the experience-dependent changes in cholinergic neuron synapses in response to forced swim, we swam the fish in a petri dish with a magnetic stir bar for 8 hours daily for 2 consecutive days; control fish were exposed to the same setup but without a stir bar (**Fig. 6a**). We found that this forced swim paradigm increased brevican intensity (**Fig. 6b,c**) as well as the number of Chat-PSD95^FingR^ excitatory synapses (**Fig. 6d,e**) relative to non-swum controls. To define synapse dynamics we adapted our time-lapse imaging paradigm to image the same neurons before and after forced swim (two sessions 30 hours apart, **Fig. 6f**). We found that forced swim increased the number of newly observed synapses detected at t=30, without altering the number of stable synapses (**Fig. 6g,h, Extended Data Fig. 10a**). Taken together, these data suggest that the forced swim experience drives concordant increases in both brevican density and excitatory synapses, driven by an increase in newly stabilized newborn synapses.

**Fig. 6:**
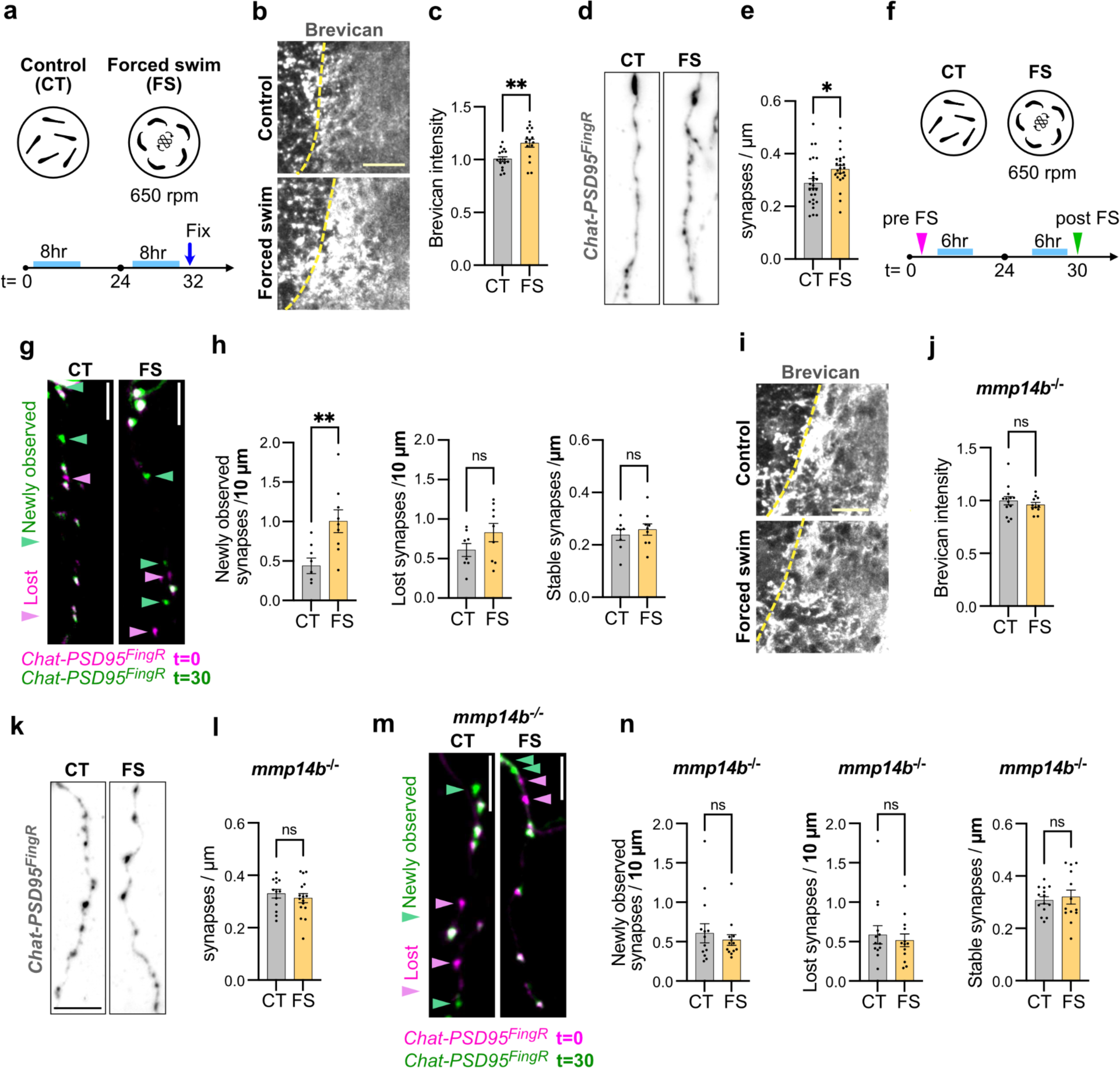
Extracellular matrix accumulation blunted experience-dependent synaptic change. a) Schematic of the forced swim paradigm performed in this study. Forced swim (FS) groups were put on a petri dish with a stirrer bar and exposed to water current for 8 hours per day at 650 rpm, for two consecutive days. Control (CT) groups were put in a petri dish without stirrer bars. Experiments were performed at 13-14 dpf. b) Representative images of brevican staining in control and forced swim groups. Scale: 20 µm. c) Quantification of brevican intensity in synaptic region normalized to mean of control group (CT, n=16 fish; FS, n=17 fish; p=0.0021**, Welch’s t-test). d) Representative images of *Chat-PSD95^FingR^* dendrites from fixed sections stained for TdT in forced swim or control group. Scale: 5 µm. e) Quantification of excitatory synapse density (Chat-PSD95^FingR^ puncta) per µm of dendrite length in forced swim or control group. Dots represent means per fish from at least one dendritic segment analyzed per fish (CT, n=21 fish; FS, n=21 fish; p=0.026*, Welch’s t-test). f) Schematic of time lapse synapse imaging performed before (t=0) and after (t=30) forced swim paradigm. Experiments were performed at 10-12 dpf. g) Representative merged images of time lapse assay in the *Chat-PSD95^FingR^* before (t=0, pink) and after (t=30, green) forced swim paradigm or control. Pink arrowheads: lost synapses, green arrowheads: new synapses. Non-merge images in Extended Data Fig.10a. Scale: 5 µm. h) Quantification of newly observed, lost, and stable synapses after forced swim paradigm (CT, n=8 fish; FS, n=9 fish; p=0.0060** for newly observed, p=0.140 for lost, p=0.507 for stable, Welch’s t-test). i) Representative images of brevican staining in *mmp14b^-/-^* fish from forced swim or control group. Scale: 20 µm. j) Quantification of brevican intensity in synaptic region in *mmp14b^-/-^* fish normalized to control group (CT, n=13 fish; FS, n=11 fish; p=0.397, Welch’s t-test). k) Representative images of *Chat-PSD95^FingR^* dendrite in *mmp14b^-/-^* fish from forced swim or control group. Scale: 5 µm. l) Quantification of synapse density (Chat-PSD95^FingR^ puncta) per µm of dendrite length from forced swim or control group in *mmp14b^-/-^*. Dots represent means per fish from at least 1 dendritic segment analyzed per fish (CT, n=13 fish; FS, n=16 fish; p=0.48, Welch’s t-test). m) Representative merged images of time lapse assay in the *Chat-PSD95^FingR^;mmp14b^-/-^* fish before (t=0, pink) and after (t=30, green) forced swim paradigm or control. Pink arrowheads: lost synapses, green arrowheads: new synapses. Non-merged images in Extended Data Fig. 10b. Scale: 5 µm. n) Quantification of newly observed, lost, stable synapses after forced swim in the *mmp14b^-/-^* fish (CT, n=14 fish; FS, n=13 fish; p=0.542 for newly observed, p=0.621 for lost, p=0.690 for stable, Welch’s t-test). Values were plotted as mean ±SEM. **: p<0.01; *: p<0.05; ns: not significant.

We next examined the effect of forced swim in fish that lacked Mmp14b. Here we found that brevican density failed to increase in response to forced swim (**Fig. 6i,j**), and excitatory synapses also did not increase, although they were higher at baseline than *mmp14b^+/+^* as previously shown (**Fig. 6k,l**). Time-lapse imaging of synapse dynamics (**Fig. 6m,n, Extended Data Fig. 10b**) also did not reveal the increase in newly observed synapses in *mmp14b^+/+^*fish. Finally, we also failed to see an increase in excitatory synapses in *bcan^-/-^*fish (**Extended Data Fig. 10c**). Taken together, these results suggest that both stable amounts of ECM (brevican) and the capacity for ECM remodeling are needed to permit new excitatory synapse formation in this model of experience-dependent neural circuit plasticity.

### Computational modeling supported the existence of two synapse states and a role of ECM in stabilizing newborn synapses

Our data on synapse dynamics using ECM loss and gain of function perturbations suggest that ECM stabilizes newborn synapses thereby increasing synapse number during development. However, as we cannot monitor synapses continuously, our data is limited by the experimentally determined imaging windows. We therefore sought to develop a mathematical model of developmental synapse dynamics that fit our experimental observations and could potentially predict the rate at which synapses are formed and remodeled in control and ECM-altered conditions.

Based on our experimental observations, we considered a model whereby developing synapses can exist in two states: a newborn unstable state and a stable state in which synapse survival is higher (**Fig. 7a**, see **Extended File 1** for mathematical derivations). Four primary parameters define the behavior of this model: (1) the synapse birth rate (b_n_), (2) the probability that a synapse will convert from the unstable to stable state (c_n->s_), (3) the decay rate of the newborn state (d_n_), and (4) the decay rate of the stable state (d_s_). Regarding the birth rate, we assume that new synapses are generated at a certain rate via a homogeneous poisson process. We further assume that a newly born synapse will transition to a stable state or decay at a rate that is proportional to its birth rate, and that once in a stable state, synapses may still decay but at a slower rate. The total number of new and stable synapses are thus random variables that evolve according to the above stochastic dynamics and can be predicted using our model.

**Figure 7:**
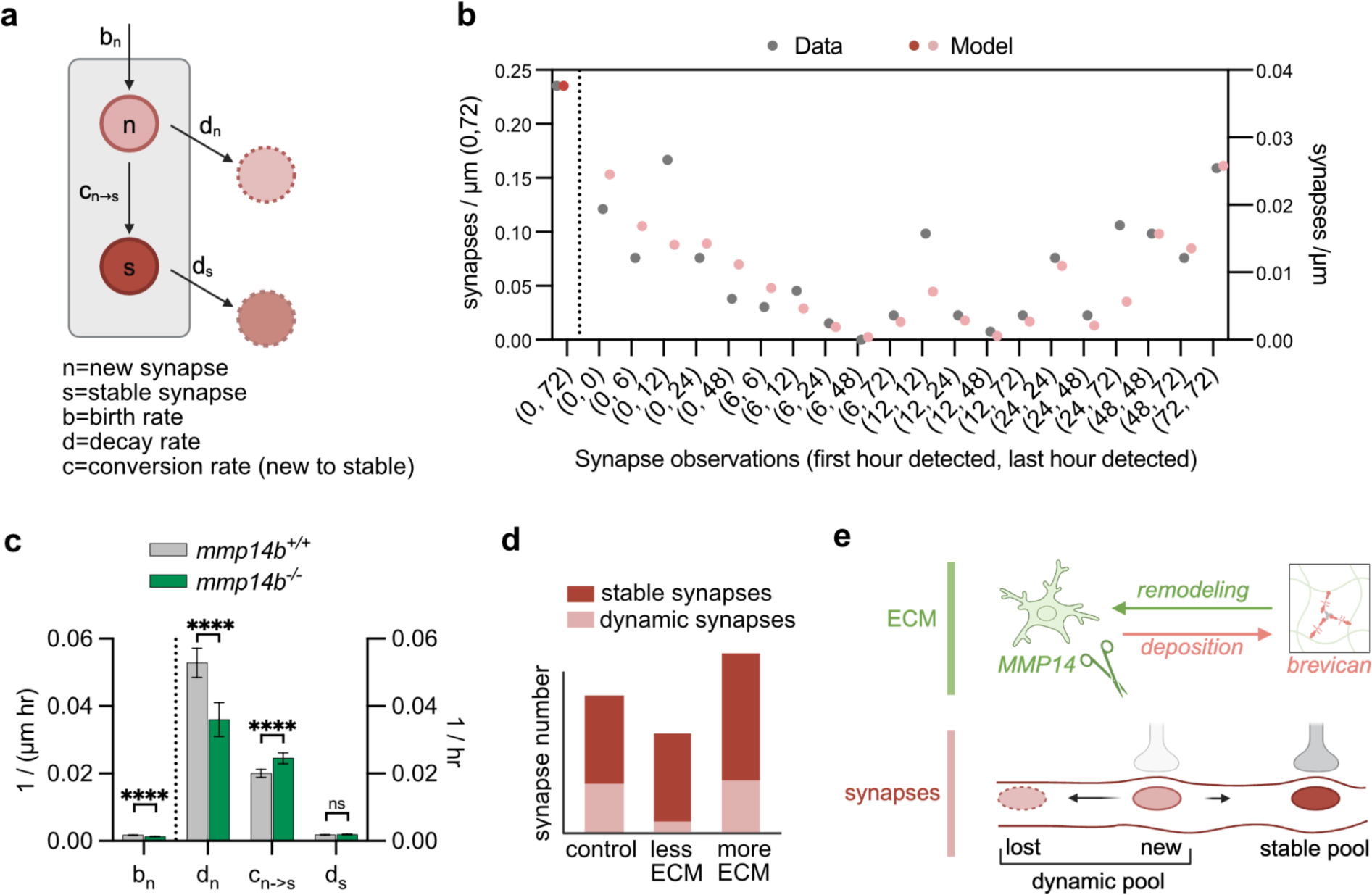
Computational modeling supported the existence of two synapse states and a role of ECM in stabilizing newborn synapses. a) Model of synapse dynamics in our system, based on four parameters: birth rate of new synapses (b_n_), decay rates of new synapses (d_n_), probability of transition from new to stable synapse (c_n->s_), and decay rate of stable (d_s_) synapses. b) Fitting of synapse lifetime data in Fig. 1m from *Chat-PSD95^FingR^* fish, based on the two state model described in panel a. Density of synapses for each category of lifetime are shown from raw data (gray) and as predicted by the two-state model (red). The numbers in parentheses on the x-axis indicate the first and last time (in hours) that synapses were detected over 72 hours. Note that while experimental analyses in Figures 1 and 3 tracked only synapses that were present at 0 hours (“stable”) or born between 0 and 6 hours (“new”), the computational model examined all synapses observed over 72 hours of imaging to generate this predicted distribution. c) Synapse dynamics calculated from the two-state model in *mmp14^+/+^* and *mmp14b^-/-^* fish, showing calculations of the four parameters described in panel a. Units for synapse birth rate (b_n_) reflect synapse born / µm / hour shortened to 1 / (µm hr). Birth rate is independent of the number of synapses in the system. Units for rates of synapse decay or stabilization (d_n_, c_n->s_, and d_s_) reflect the rates as a proportion of existing synapses, i.e. changed synapses / µm / hour divided by total synapses / µm, shortened to 1/ hr. Error bars show ±SD. ****: p<0.0001, Mann Whitney U test. d) Graphical data summary. e) Graphical abstract of ECM impact on synapse numbers and dynamics.

To test how this two-state model aligned with our experimentally determined data, we used the model to predict synaptic observations made during a 72 hours imaging period and compared this model against our experimentally determined synapse lifetime data from Figure 1k-m. We found that the model accurately predicted the ratio of dynamic to stable synapses observed experimentally in control data from Figure 1m (**Fig. 7b**) and in data from *mmp14b^-/-^*fish in Fig. 3q (**Extended Data Fig. 11a**). In contrast, a single-state model of synapse dynamics was unable to fit the experimentally observed data (**Extended Data Fig. 11b,c**).

We next used this model to calculate synapse birth rate, decay rates, and conversion rates from dynamic to stable in control and *mmp14b* deficient conditions – parameters can be extrapolated from our model but are challenging to accurately determine experimentally (**Fig. 7c**). Examining only our control data, our model predicted a synapse birth rate of 0.0017 synapses/μm/hour. It also predicted that the decay rate of newborn synapses is much larger than the decay rate of stable synapses, as expected (d_n_=0.053 vs. d_s_=0.002 per hour). It further predicts that a majority of new synapses will decay, rather than be converted to stable synapses (d_n_=0.053/hr vs. c_n->s_ = 0.02/hr). In the setting of *mmp14b* deficiency, the model predicted a 26% reduction in the synapse birth rate suggesting that a change in birth rate could not explain the increased number of “newly observed” synapses seen over a 24 hour imaging window (see Fig. 3n). However, the decay rate of new synapses (d_n_) was decreased by 32%, and the transition of new to stable synapses (c_n->s_) was increased by 22%. Taken together, these computations supported our hypothesis that remodeling of the ECM maintains synapses in a dynamic state and that impaired ECM remodeling leads to an accumulation of stable synapses (**Fig. 7d,e**). This would be predicted to diminish the efficiency of experience dependent adaptation and could contribute to the observed circuit and behavioral impacts of ECM alterations.

## Discussion

Our study directly examined how alterations of the brain ECM shape the structural plasticity of developing synapses and identified brevican and microglial MMP14 as a molecular regulator of this process. We propose a model whereby MMP14 acts by promoting microglial cleavage of brevican, which destabilizes and preferentially eliminates recently born synapses. In support of this model, we show that loss of either microglia or MMP14 in developing fish leads to accumulation of brevican, an increase in synapses, and an increase in newly stabilized synapses. Conversely, loss of brevican leads to a loss of synapses, consistent with studies in mice^18^, and a decrease in newly stabilized synapses. We show both in fish and in human tri-cultures that microglial MMP14 mediates this effect. It is possible that MMP14 acts on additional ECM proteins besides brevican, or has alternate functions besides ECM cleavage.

For example, the behavioral deficits in brevican deficient fish are not just the converse of MMP14 deficiency. While this could be explained by the total loss of brevican in the knockout, which is more profound than the 20% increase seen in MMP14 deficient mice, it is also likely that additional ECM proteins are cleaved by MMP14^69^.

Our data suggest a novel mechanism through which microglia promote developmental synapse elimination: by destabilizing newborn synapses and promoting their subsequent loss via neuron autonomous mechanisms. Many studies suggest that molecular pathways that promote microglial function also restrict synapse numbers in the developing brain^70–72^. While this is often interpreted as evidence of microglial synapse pruning, or engulfment, a close review of the literature has not identified instances of microglia directly engulfing synapses *in vivo*^73,74^. This mechanism is distinct from a prevailing model whereby microglia prune synapses by active synaptic engulfment. In this alternate model, we propose that microglia alter the extracellular environment of synapses in a manner that promotes their subsequent elimination or resorption by the neuron.

It is possible that microglial ECM alterations occur in a spatially restricted manner. For example, we observed that microglial processes frequently contacted excitatory synapses, similar to mice^36^. It is interesting to speculate that microglial cleavage of the ECM could be contact-dependent as MMP14 is surface bound and was concentrated in microglial processes. Interestingly, inhibition of another membrane-bound metalloproteinase Adam10 in microglia prevented synapse elimination after whisker deprivation in developing mouse sensory cortex^75^. However, support for our model of microglial contact-dependent synapse elimination by ECM clearance would require further experiments to determine (1) whether synapses previously contacted by microglia are eventually eliminated, and (2) whether this effect depends on the amount of ECM in the local environment. Advances in non-invasive microscopy such as lattice light sheet imaging could enable prolonged live imaging of structural synaptic changes, which can occur on the order of hours to days^51^, and ideally, concurrent imaging of the ECM itself. Whether synapse specific or not, microglial remodeling of the ECM could explain many scenarios in which microglia contribute to synapse elimination.

These findings also raise a critical question: if microglial remodeling of the ECM in development leads to synapse elimination, why does it have the opposite effect of promoting synapse formation in adulthood, as we and others have previously shown?^29,76,77^ This is not simply a difference in model systems, as we showed here that two independent methods of ECM depletion reduced synapse numbers in development but increased them in adulthood in the same circuit and synapse type. The answer to this question may reflect both developmental differences in the ECM, and differences in the structural plasticity of synapses. With regards to the ECM, we and others have shown a rapid accumulation of ECM proteins like brevican in the early postnatal period. These proteins reach maximal levels and change their glycosylation patterns by adulthood^17,43^. ECM can both stabilize synapses and prevent synaptic plasticity. In development, a relatively looser ECM may stabilize newborn synapses but be easily displaced as new synapses are formed. In adulthood, a denser ECM lattice may form a steric barrier to new synapse formation. Further studies and computational modeling of how ECM function changes with age could extend these findings to other contexts, such as to explain the relationship between ECM accumulation and the closure of developmental critical periods.

A second fundamental difference between development and adulthood is the higher fraction of developing synapses that undergo structural plasticity. While synapse and synaptic protein turnover persists well into adulthood^78^, the fraction of synapses that are dynamic decrease^52^. Our data suggests that most excitatory synapses are short-lived in developing cholinergic neurons in zebrafish hindbrain, consistent with other studies^39^. Increased ECM preferentially increased the number of these nascent synapses, while having relatively little effect on the stable synapse pool. Our data suggest that similar to other synaptic organizers such as neurexins^79^, this effect of the ECM reflects increased stabilization of newborn synapses rather than a direct induction of new synapse formation. Therefore, loss of ECM would preferentially reduce overall synapse numbers in developing brains despite increasing synaptic plasticity in both settings. As zebrafish lose optical transparency around 14 dpf, live imaging in other model organisms (such as the related fish species *Danionella*^80^) could directly test this hypothesis in the same circuit and brain region.

Finally, structural synaptic plasticity is required for experience-dependent adaptation both during development and in adulthood. Our data suggest that ECM remodeling is essential for this plasticity. We found that forced swim, a stressor and inducer of motor learning, led to a robust increase in newborn synapses, consistent with studies of motor learning in mouse cortical neurons^52^. This increase was completely abrogated in MMP14 deficient fish. Chronic stress models in rodents cause an increase of brevican and decrease of MMPs^81,82^, suggesting that both ECM deposition and remodeling could be altered. Thus ECM remodeling could be essential to optimize circuit function in response to stress and novel experience. Given that MMP14 is highly expressed in human microglia, this mechanism of ECM remodeling may be conserved in the human brain and may suggest new strategies to regulate plasticity in human neurodevelopmental disorders.

## Acknowledgements

We are grateful to Molofsky lab members for helpful feedback and support. We are also grateful to the UCSF zebrafish core facility for help in zebrafish husbandry, and the UCSF Center for Advanced Light Microscopy for help in imaging. Thanks to Dr. Francesca Peri for the *Tg(mpeg1.1:GFP-CAAX)* fish, Dr. Brian Black for the *mmp14b^-/-^* fish, Dr. Fumi Kubo for the *Tg(chata:gal4)* fish, Dr. Joshua Bonkowsky for the FingR construct, Dr. Kelly Smith for the *Tg(ubi:ssncan-GFP)* fish, Dr. Kelly Monk for the *Tg(slc1a3b:myrGFP-P2A-H2A-mCherry)* fish, Dr. Helen Willsey and Dr. Scott Baraban for use of the DanioVision system, Dr. John Boscardin for statistics consultation, and Kunal Shroff and Olivia Teter for experimental guidance. We acknowledge the following funding support: to A.V.M.: DP2MH116507, the Brain Research Foundation, The Program for Breakthrough Biomedical Sciences (PBBR); to H.N.: Astellas Foundation for Research on Metabolic Disorders, The Uehara Memorial Foundation, Japan Society for the Promotion of Science (JSPS); to D.L.S.: NIH R01AG085357 and U54NS123746.

## Author contributions

Conceptualization, H.N., and A.V.M.; methodology, H.N., and R.C.; investigation, H.N., R.C., S.A.M., N.J.S., M.S., K.S., I.R., J.M.; computational modeling, C.K.; writing-original draft, H.N., and A.V.M.; writing-review & editing, all co-authors; funding acquisition, A.V.M.; D.L.S.; resources, A.V.M.; supervision, A.V.M.; D.L.S., M.K., C.K.

---

**Extended Data Figure 1:**
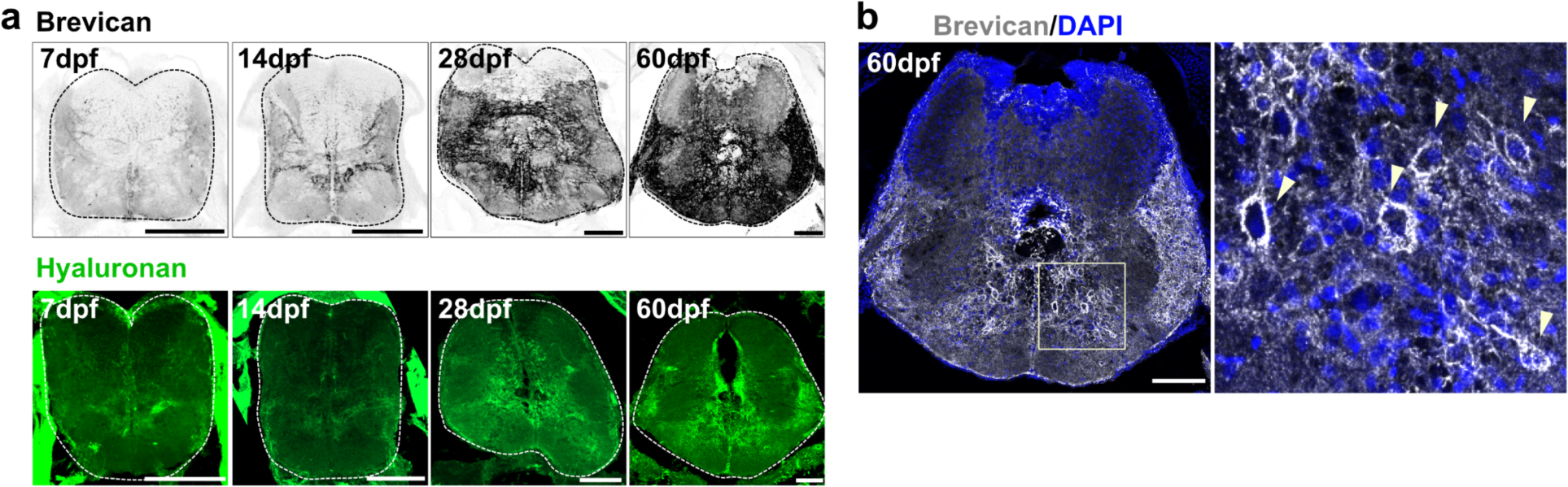
Characterization of ECM over development in hindbrain. a) Representative images of brevican and hyaluronan (*ubi:ssncan-GFP*) in the hindbrain at 7, 14, 28, and 60 dpf. Scales: 100 µm. b) Representative image of PNN (perineuronal net)-like brevican staining at 60 dpf. Square indicates the region of inset on the right. Arrowheads indicate PNN-like brevican signals. Scale: 100 µm.

**Extended Data Fig. 2:**
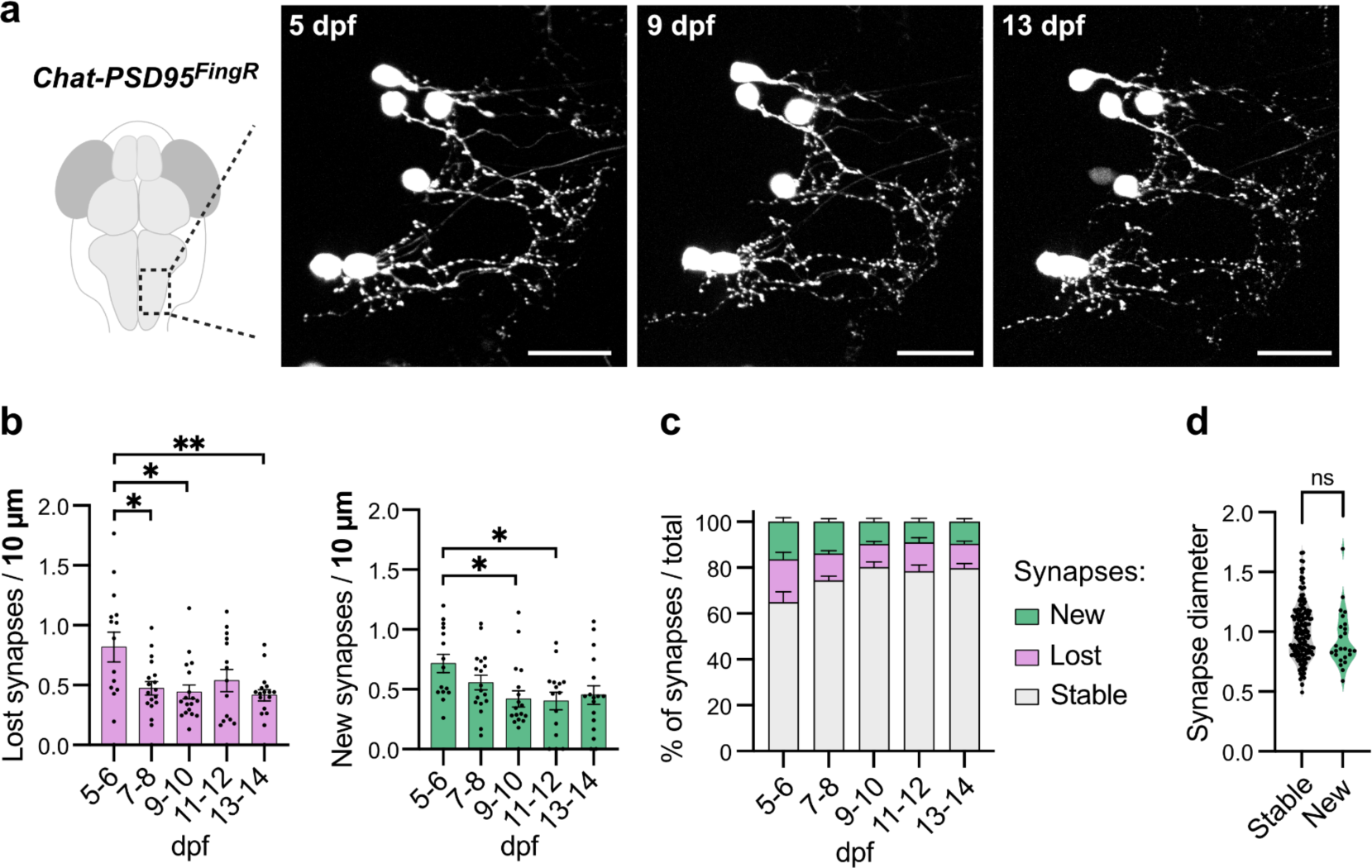
Further assessment of synapse dynamics in hindbrain cholinergic neurons. a) Representative images of *Chat-PSD95^FingR^*at 5, 9, and 13 dpf from the hindbrain of the same fish. Scales: 20 µm. b) Plots of lost (left, pink) and new (right, green) synapses for individual fish from Fig. 1j. Individual dots indicate the value for each fish (5-6 dpf, n=15 fish; 7-8 dpf, n=18 fish; 9-10 dpf, n=18 fish; 11-12 dpf, n=15 fish; 13-14 dpf, n=17 fish). Asterisks in the figure represent results of Tukey’s multiple comparisons with respect to the 5-6 dpf shown in Fig. 1j.. c) Quantification of percentage of new, lost, and stable synapses in each measured time point of Fig. 1j. Total number of synapses were calculated as the sum of the number of newly observed, lost and stable synapses. d) Quantification of synapse diameter from stable synapses and new synapses at t=6 (Stable, n=157 synapses from n=14 fish; New, n=25 synapses from n=14 fish; p=0.206, Welch’s t-test, performed for synapses). The synapse diameters were normalized by the mean value of stable synapses for each cell. Values were plotted as mean ±SEM. ns: not significant.

**Extended Data Figure 3:**
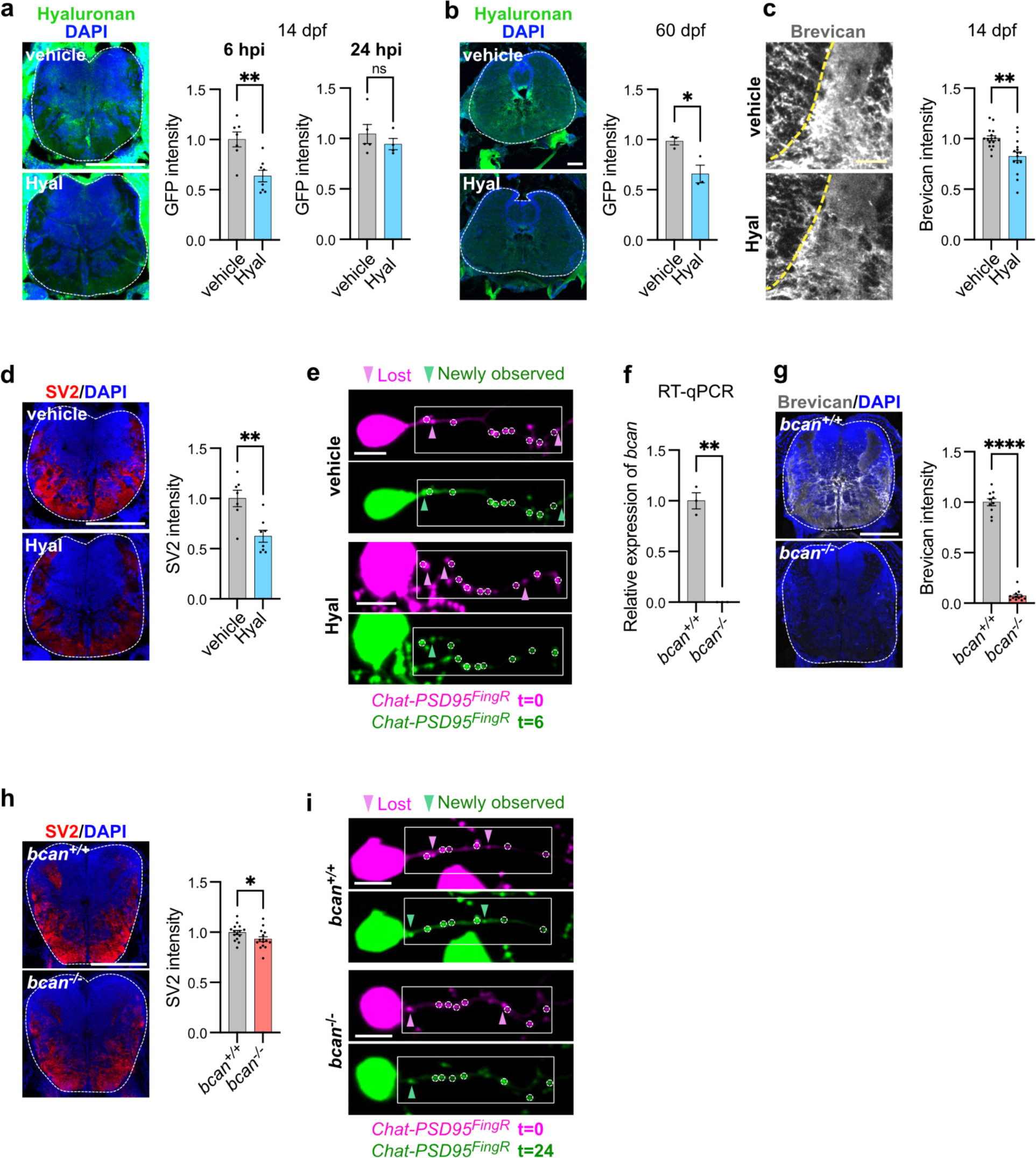
Optimization and validation of ECM digestion and *bcan* deletion. a) Representative images and quantification of hyaluronan depletion after hyaluronidase injection at 14 dpf, measured as mean fluorescence intensity of GFP (*ubi:ssncan-GFP*) at 6 hours post injection (hpi) (vehicle, n=7 fish; Hyal, n=8 fish; p=0.0023**, Welch’s t-test) and 24 hpi (vehicle, n=5 fish; Hyal, n=4 fish; p=0.392, Welch’s t-test). GFP intensity normalized to mean of vehicle control. Dashed lines indicate hindbrain regions. Scale: 100 µm. b) Representative images and quantification of hyaluronan depletion at 6 hpi at 60 dpf, measured as mean fluorescence intensity of GFP (*ubi:ssncan-GFP*) nomalized to vehicle control (vehicle, n=3 fish; Hyal, n=3 fish; p=0.049*, Welch’s t-test). Scale: 100 µm. c) Representative images and quantification of brevican depletion at 6 hours after hyaluronidase injection at 14 dpf (vehicle, n=19 fish; Hyal, n=18 fish; p=0.0025**, Welch’s t-test). Brevican intensity normalized to mean of vehicle control. Dashed lines indicate hindbrain regions. Scale: 20 µm. d) Representative images and quantification of SV2 presynapse marker 6 hours after hyaluronidase injection at 14 dpf (vehicle, n=7 fish; Hyal, n=8 fish; p=0.003**, Welch’s t-test). SV2 intensity normalized to mean of vehicle control. Scale: 100 µm. e) Non-merged images of time lapse imaging from Fig. 2d. Circles indicate stable synapses. Boxes indicate the region shown in Fig. 2d. Scale: 5 µm. f) Expression of *bcan* gene in *bcan^-/-^*fish assessed by RT-qPCR from whole larvae. Expression normalized to *ef1a* housekeeper gene (n=3/group; p=0.0064**, Welch’s t-test). g) Representative images and quantification of brevican protein in *bcan^-/-^* fish (*bcan^+/+^*, n=10 fish; *bcan^-/-^*, n=12 fish; p<0.0001****, Welch’s t-test). Brevican intensity normalized to mean of *bcan^+/+^* control. Scale: 100 µm. h) Representative images and quantification of SV2 presynapse marker in *bcan^-/-^* fish at 14 dpf (*bcan^+/+^*, n=15 fish; *bcan^-/-^*, n=15 fish, p=0.049*, Welch’s t-test). SV2 intensity normalized to mean of *bcan^+/+^*control. Scale: 100 µm. i) Non-merged images of time lapse imaging experiment from Fig. 2i. Circles indicate stable synapses. Boxes indicate the region shown in Fig. 2i. Scale: 5 µm. Values were plotted as mean ±SEM. ****: p<0.0001; **: p<0.01; *: p<0.05; ns: not significant.

**Extended Data Figure 4 :**
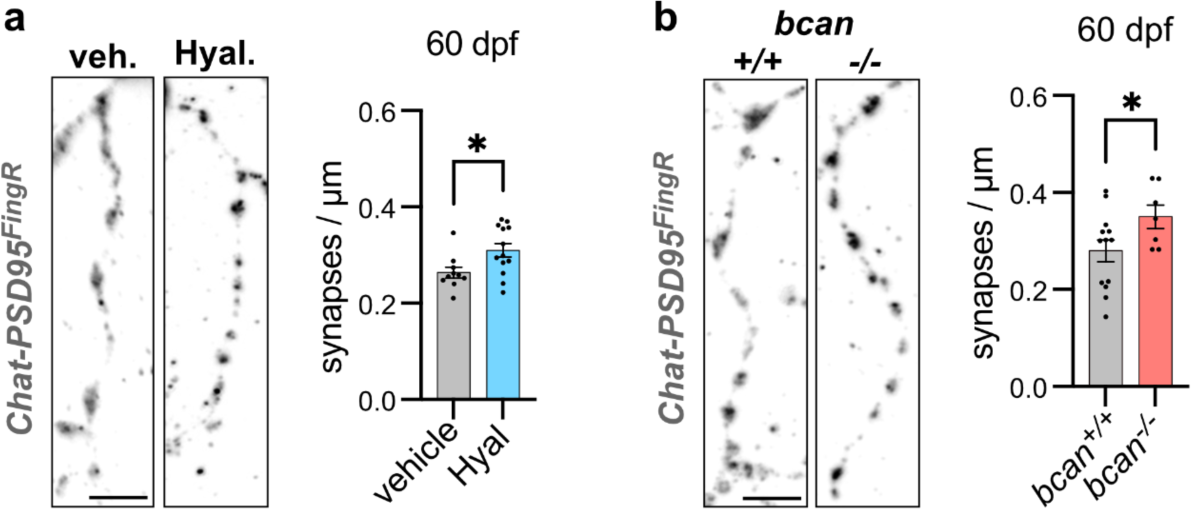
ECM depletion increases synapse density in the adult brain. a) Representative images and quantification of *Chat-PSD95^FingR^* at 60 dpf from fixed section stained for ΤdT six hours after vehicle (PBS) or hyaluronidase injection. Dots represent means per fish, with at least 1 dendrite quantified per fish (vehicle, n=10 fish; Hyal, n=13 fish; p=0.018*, Welch’s t-test). Scale: 5 µm. b) Representative images and quantification of *Chat-PSD95^FingR^* dendrites at 60 dpf from fixed section stained for ΤdT from *bcan^+/+^*vs. *bcan^-/-^* fish. Dots represent means per fish, with at least 1 dendrite quantified per fish (*bcan^+/+^*, n=13, *bcan^-/-^*, n=7; p=0.048*, Welch’s t-test). Scale: 5 µm. Values were plotted as mean ±SEM. *: p<0.05.

**Extended Data Fig. 5:**
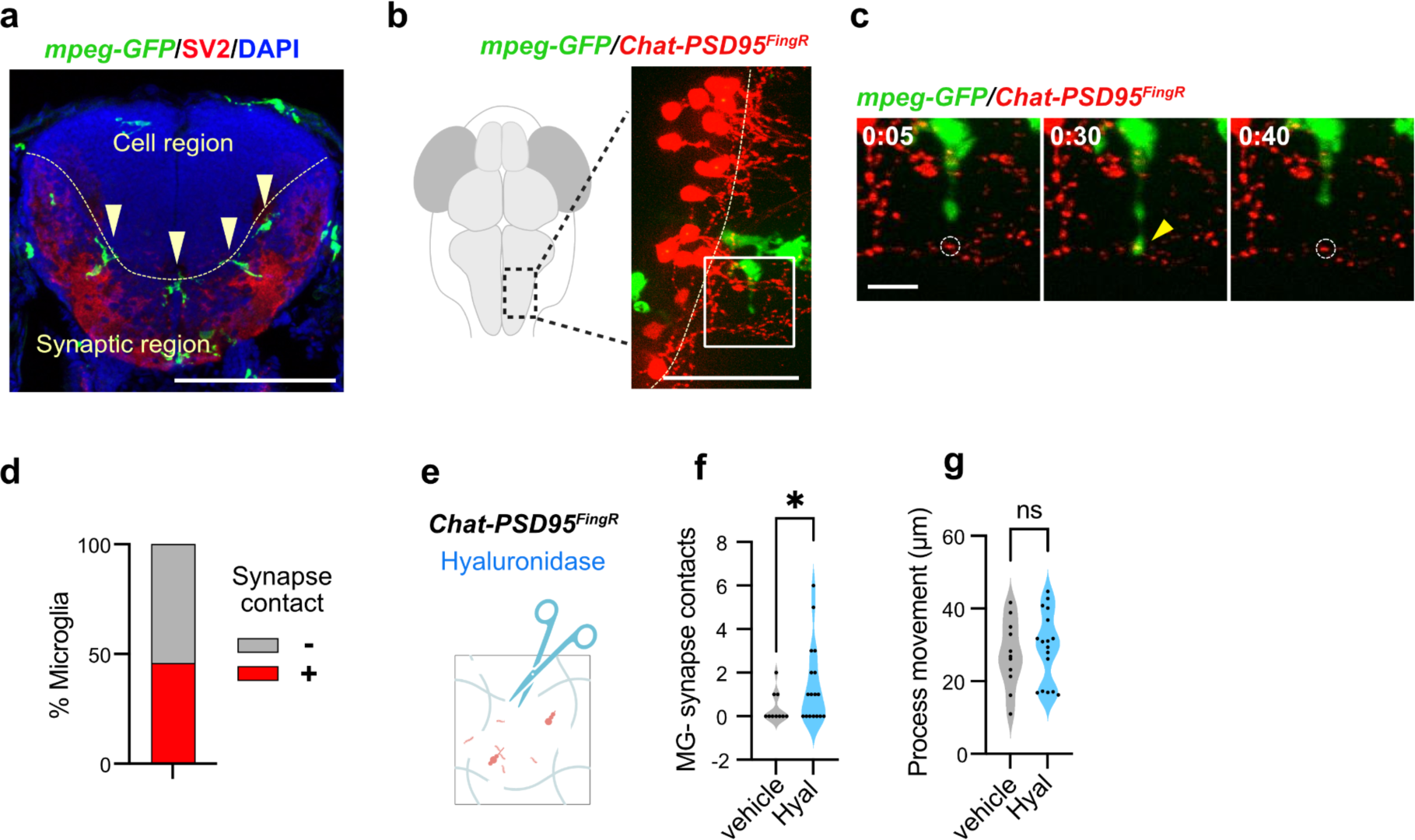
Microglia contact excitatory synapses. a) Representative image of microglia and SV2 presynaptic marker in the coronal hindbrain section from *mpeg-GFP* fish at 14 dpf. The DAPI-rich region (cell region) and synapse-rich region (synaptic region) are indicated in the image. Arrowheads indicated microglia associated with the synaptic region. Scale bar, 100 µm. b) Schematic of live imaging shows a dorsal view of hindbrain with microglia labeled with *mpeg-GFP,* and sparsely labeled cholinergic neuron and synapses with *Chat-PSD95^FingR^*. Scale: 50 µm. c) Representative time lapse image of microglia contact with *Chat-PSD95^FingR^* puncta. Z-stack images taken every 5 min for 1 hr. Dashed circles indicate *Chat-PSD95^FingR^* puncta before and after microglial contact and yellow arrowhead indicates contact detected by colocalization of TdT and GFP. Scale: 10 µm. See also Supplementary video 1. d) Quantification of percentage of microglia that contacted *Chat-PSD95^FingR^* puncta during 1 hr video (n=24 microglia from 13 fish). e) Schematic of hyaluronidase injection to digest ECM before live imaging of microglial-synapse contacts. f) Quantification of the number of microglial contacts with *Chat-PSD95^FingR^* puncta during 1 hr video (vehicle, n=10 microglia from 5 fish; Hyal, n=17 microglia from 8 fish; p=0.043*, Welch’s t-test, performed for microglia). g) Quantification of microglial process movement during 1 hr video (vehicle, n=11 microglia; Hyal, n=17 microglia; p=0.606, Welch’s t-test, performed for microglia). Values were plotted as mean ±SEM.*: p<0.05; ns: not significant.

**Extended Data Fig. 6:**
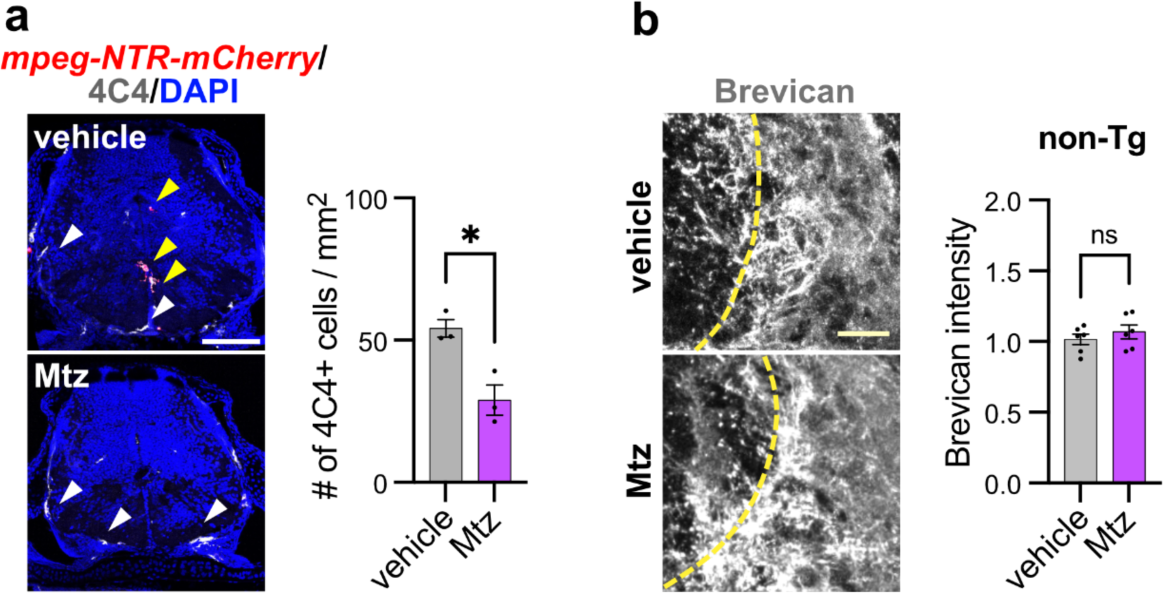
Validation of microglial depletion strategy using nitroreductase. a) Representative images of mCherry and the microglial marker 4C4 and quantification of microglia number in the *Tg(mpeg:gal4);Tg(UAS:NTR-mCherry)* with Mtz or vehicle (DMSO) treatment at 14 dpf (vehicle, n=3 fish; Mtz, n=3 fish; p=0.015*, Welch’s t-test). non-Tg= non-transgenic fish. Scale: 100 µm. b) Representative images and quantification of brevican staining in Mtz-treated non-transgenic siblings at 14 dpf (vehicle, n=6 fish; Mtz, n=6 fish; p=0.412, Welch’s t-test). Brevican intensity normalized to the mean of vehicle control. Scale: 20 µm. Values were plotted as mean ±SEM.*: p<0.05; ns: not significant.

**Extended Data Fig. 7:**
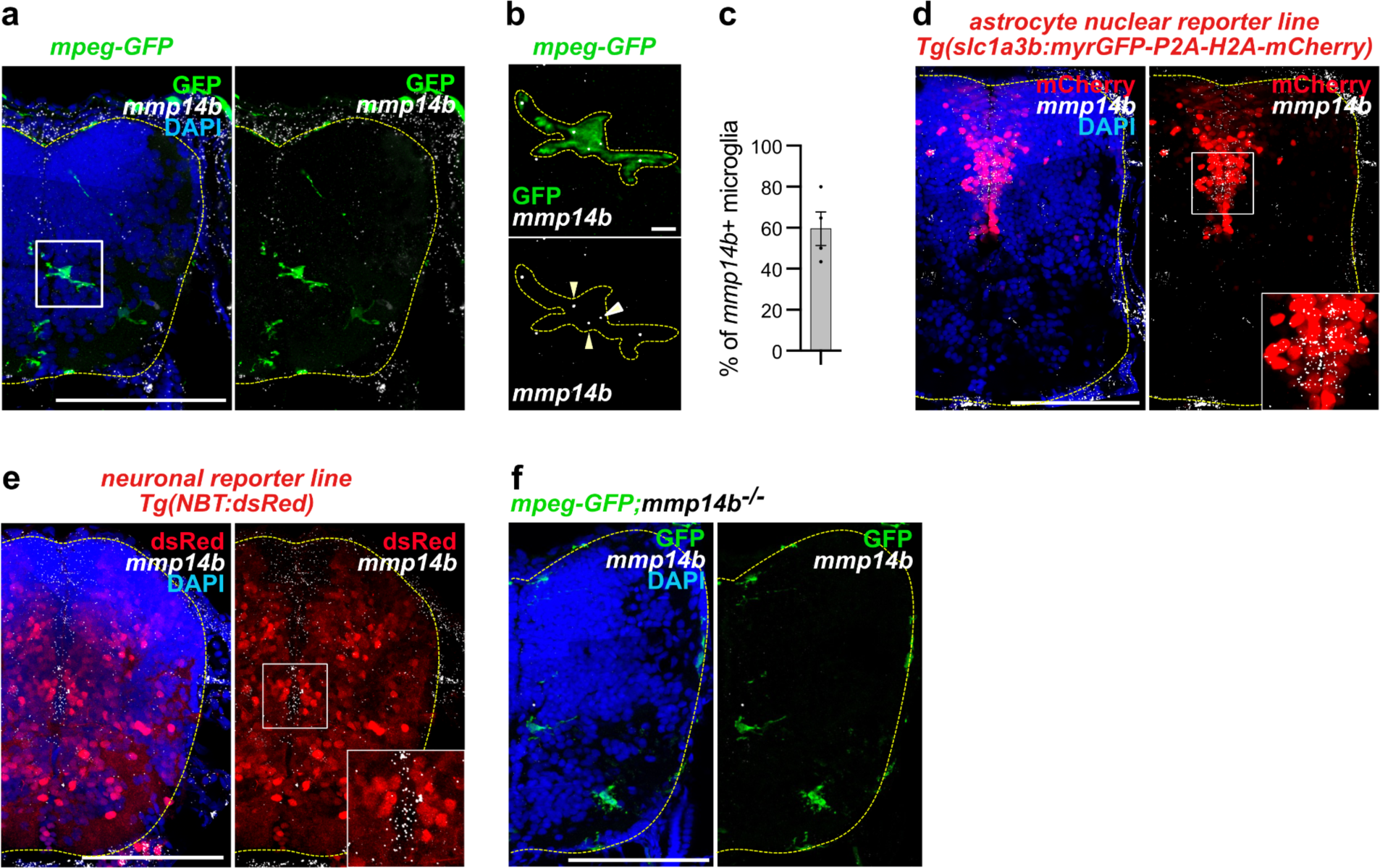
*mmp14b* is expressed in microglia and astrocytes, but not neurons. a) Representative image of RNA *in situ* hybridization for *mmp14b* in *mpeg-GFP* fish. Dashed lines indicate hindbrain regions. White box indicates inset in b. Scale: 100 µm. b) Inset of microglia in a. Arrowheads indicate *mmp14b* puncta in *mpeg-GFP* microglia. Dashed lines indicate microglia. Scale: 5 µm. c) Quantification of percentage of hindbrain microglia expressing *mmp14b.* Dots indicate n=4 fish. d) Representative images of RNA *in situ* hybridization for *mmp14b* in astrocyte reporter line *Tg(slc1a3b:myrGFP-P2A-H2A-mCherry)* at 14 dpf. The mCherry signals indicate the astrocyte nucleus. Dashed lines indicate hindbrain regions. Inset is shown at the bottom. Scale: 100 µm. e) Representative images of RNA *in situ* hybridization for *mmp14b* in neuronal reporter line *Tg(NBT:dsRed)* at 14 dpf. Dashed lines indicate hindbrain regions. Inset is shown at the bottom. Scale: 100 µm. f) Representative image of RNA *in situ* hybridization for *mmp14b* in *mpeg-GFP*;*mmp14b^-/-^* fish showing lack of probe hybridization. Dashed lines indicate hindbrain regions. Scale: 100 µm. Values were plotted as mean ±SEM.

**Extended Data Fig. 8:**
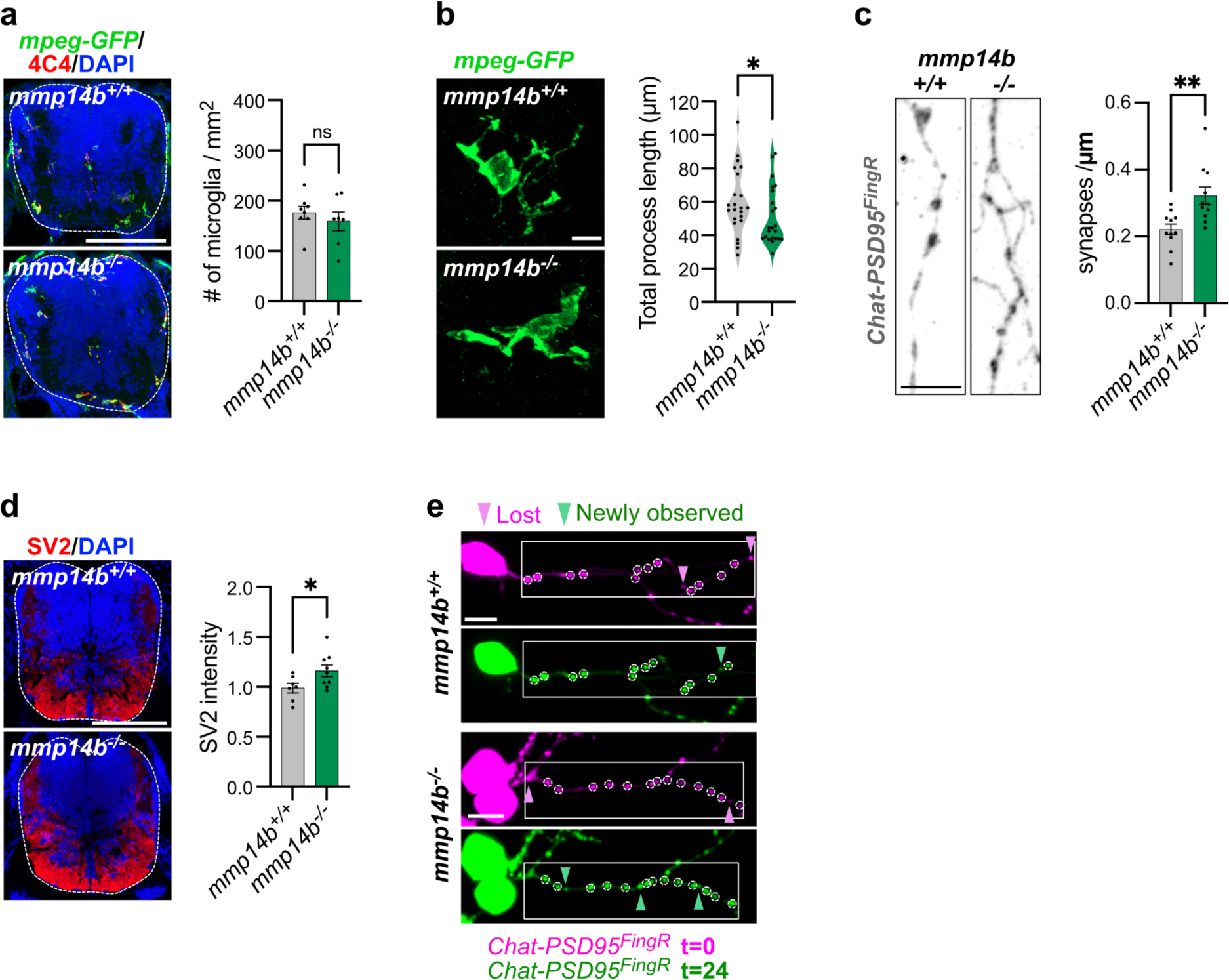
Additional characterization of *mmp14b* deficient fish. a) Representative images and quantification of the number of microglia having 4C4 microglial marker in *mpeg-GFP* fish (*mmp14b^+/+^*, n=8 fish; *mmp14b^-/-^*, n=7 fish; p=0.468, Welch’s t-test). Dashed lines indicate hindbrain regions. Scale: 100 µm. b) Representative images and quantification of the total length of microglial processes (*mmp14b^+/+^*, n=23 microglia from n=7 fish; *mmp14b^-/-^*, n=23 microglia from n=7 fish; p=0.026*, Kolmogorov-Smirnov test, performed for microglia). Scale: 5 µm. c) Representative images and quantification of synapse density per µm of dendrite length (Chat-PSD95^FingR^ puncta) in *mmp14b^-/-^* and *mmp14b^+/+^*control at 14 dpf. Dots represent means per fish from at least 1 dendritic segment analyzed per fish (*mmp14b^+/+^*, n=11 fish; *mmp14b^-/-^*, n=11 fish; p=0.0044**, Welch’s t-test). Scale: 5 µm d) Representative images and quantification of SV2 presynapse marker in *mmp14b^-/-^* fish at 14 dpf (*mmp14b^+/+^*, n=7 fish; *mmp14b^-/-^*, n=9 fish, p=0.035*, Welch’s t-test). SV2 intensity normalized to mean of *mmp14b^+/+^* control. Dashed lines indicate hindbrain regions. Scale: 100 µm. e) Non-merged images of time lapse imaging experiment from Fig. 3m. Circles indicate stable synapses. Boxes indicate the region shown in Fig. 3m. Scale: 5 µm. Values were plotted as mean ±SEM.**: p<0.01; *: p<0.05; ns: not significant.

**Extended Data Fig. 9:**
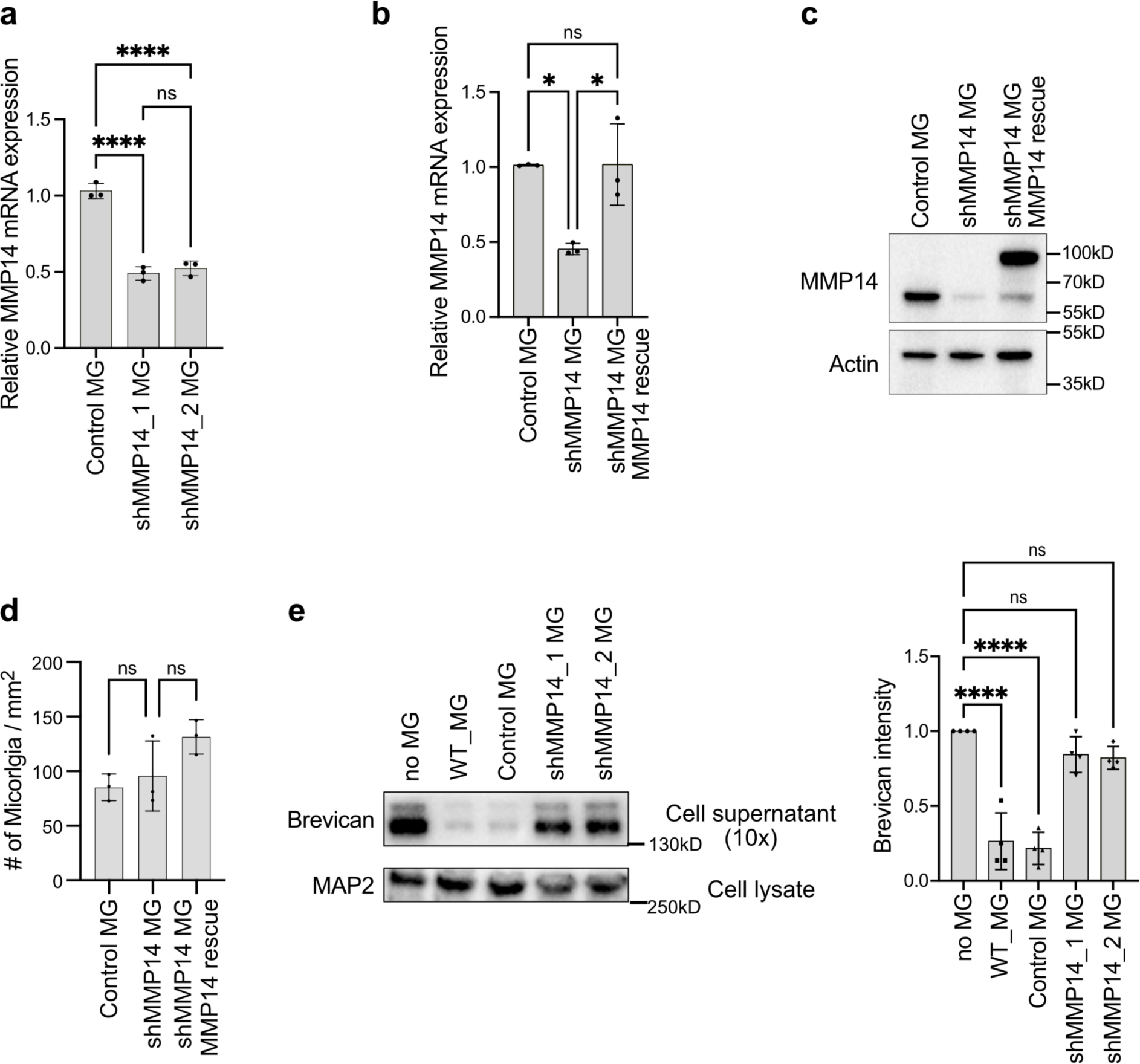
Validation of MMP14 knock-down in human iPSC derived Microglia. a) *MMP14* mRNA levels from MMP14 KD microglia analyzed by RT-qPCR showing two independent shRNAs targeting MMP14, with scrambled shRNA as control. Expression normalized to *GAPDH* housekeeping gene (n=3 independent experiments; One-way ANOVA with Tukey’s multiple comparison). b) *MMP14* mRNA levels from MMP14 KD and MMP14-GFP rescue microglia analyzed by RT-qPCR. Expression normalized to *GAPDH* (n=3 independent experiments; one-way ANOVA, asterisks in the figure show results of Tukey’s multiple comparison). c) Western blot of MMP14 showing *MMP14* KD and rescue with MMP14-GFP in microglia. Note higher molecular weight of MMP14-GFP vs. endogenous MMP14. d) Quantification of the microglia cell number in the triculture system with MMP14 KD and MMP14-GFP rescue (n=3 independent experiments, p=0.209, one-way ANOVA). e) Representative western blot and quantification of Brevican in cell supernatant vs. MAP2 loading control in cell lysate from the triculture system with MMP14 KD (n=4 independent experiments, One-way ANOVA, asterisks show results of Tukey’s multiple comparison). Values were plotted as mean ±SEM.****: p<0.0001; *: p<0.05; ns: not significant.

**Extended Data Fig. 10:**
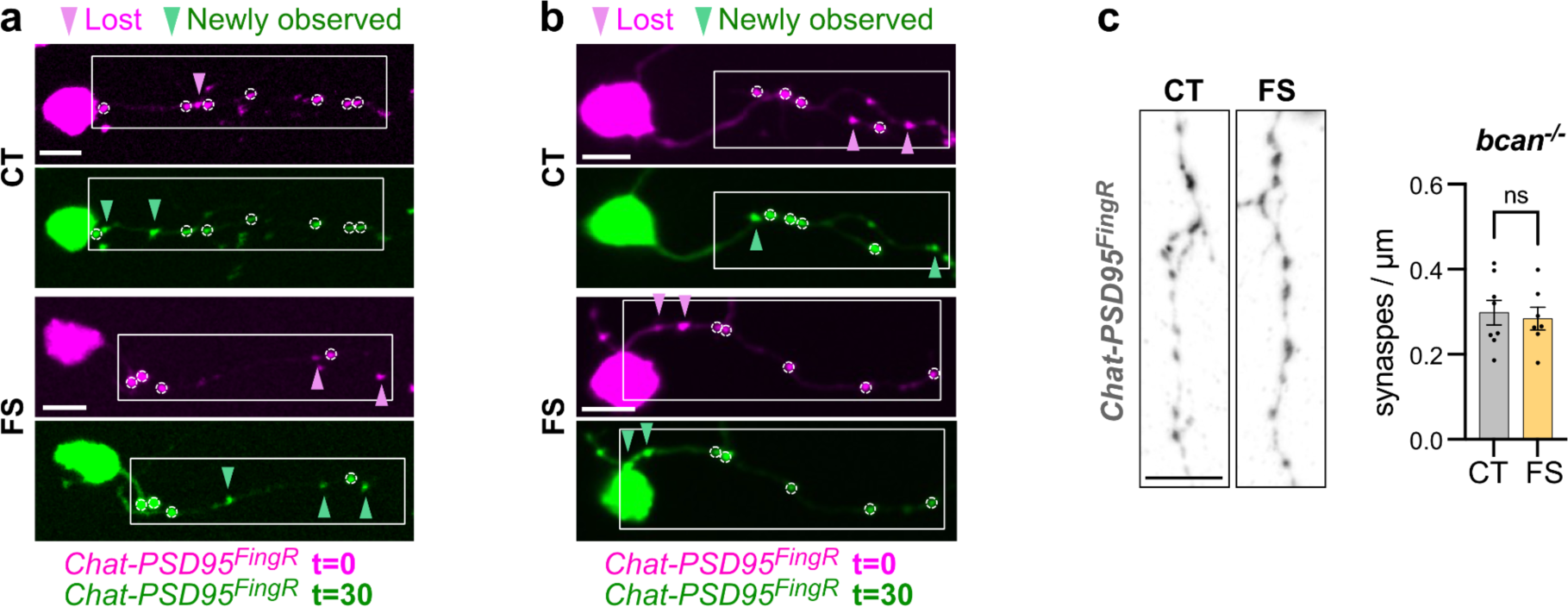
Effect of forced swim paradigm on synapses. a) Non-merged images of time lapse imaging experiment from Fig. 6g. Circles indicate stable synapses. Boxes indicate the region shown in Fig. 6g. Scale: 5 µm. b) Non-merged images of time lapse imaging experiment from Fig. 6m. Circles indicate stable synapses. Boxes indicate the region shown in Fig. 6m. Scale: 5 µm. c) Representative images and quantification of synapse density per µm of dendrite length (Chat-PSD95^FingR^ puncta) in *bcan^-/-^* fish from forced swim and control group at 14 dpf. Dots represent means per fish from at least 1 dendritic segment analyzed per fish (CT, n=8 fish; FS, n=7 fish; p=0.732, Welch’s t-test). Scale: 5 µm. Values were plotted as mean ±SEM. ns: not significant.

**Extended Data Fig. 11:**
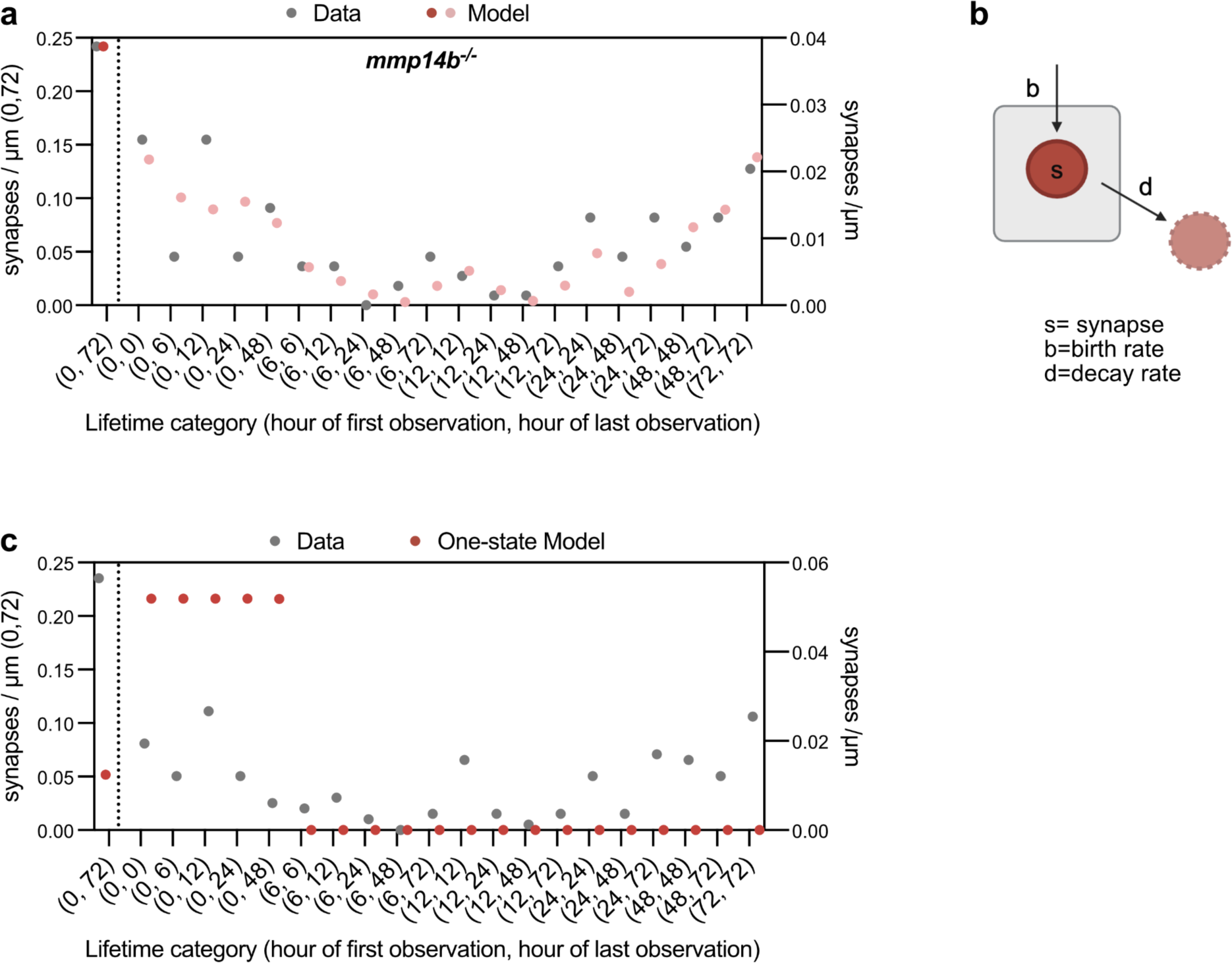
Additional computational modeling of synapse dynamics. a) Fitting of synapse lifetime data in Fig. 3q from *mmp14b^-/-^ ;Chat-PSD95^FingR^* fish, based on the two state model in Fig. 7a. Density of synapses for each category of lifetime are shown from raw data and as predicted by the two-state model. The numbers in parentheses indicate the time point of first and last observation of synapses during the time course over 72 hours. b) Schematic of a “one state” model of synapse dynamics. c) Attempted fit of experimental data from control fish in Fig 1m to a “one state” model shows poor alignment. The numbers in parentheses indicate the time point of first and last observation of synapses during time course over 72 hours.

**Supplemental Video 1**: Live imaging of microglia contact with excitatory synapses.

Representative time lapse image of microglia contact with *Chat-PSD95^FingR^* puncta shown in Extended Data Fig. 5c. Z-stack images taken every 5 min for 1 hr. White arrowheads indicate a synapse before and after microglial contact and yellow arrowheads indicate a synapse contacted by microglia.

**Supplemental Table 1**: Proteomics data used for plot in Fig. 4e.

Analyzed proteomics result used for the plot shown in Fig. 4e. Protein names, log2FC, adjusted p-values, and matrisome results are included.

**Supplemental Table 2**: Analyzed proteomics result in Fig. 4.

Analyzed expression of each protein from proteomics analysis in iPCS-derived culture supernatant. Results for 4 independent samples are included.

**Supplemental File 1**: Derivation and Fitting of the computational model.

## Materials and Methods

### Zebrafish

All animal protocols were approved by and in accordance with the ethical guidelines established by the UCSF Institutional Animal Care and Use Committee and Laboratory Animal Resource Center (LARC). Zebrafish (*Danio rerio*) were propagated, maintained, and housed in recirculating habitats at 28.5 °C and on a 14/10-h light/dark cycle. Embryos were collected after natural spawns and incubated at 28.5 °C. Sex was not considered as a variable before 28 days post fertilization (dpf) as this prior to sex determination. Adults of mixed sex were used at 60 dpf and 90 dpf of age. Ages were matched within experiments. All experiments were performed with fish on the non-pigmented Casper background (*roy^-/-^;nacre^-/-^*)^84^ except for behavior analysis, which were performed on the normally pigmented Ekkwill (EKW; ZFIN ID: ZDB-GENO-990520-2) background to preserve normal light startle responses.

### Transgenic zebrafish

The following transgenic (Tg) fish described in previous studies were used in this study ^42,47,85–87^: *Tg(mpeg1.1:GFP-CAAX)*^zh901Tg^, *Tg(mpeg1.1:gal4)*^zf2055Tg^, *Tg(UAS:NTR-mCherry)*^c264tg^, *Tg(chata:gal4)*^mpn202Tg^, *Tg(ubi:ssncan-GFP)*^uq25bhTg^, *Tg(slc1a3b:myrGFP-P2A-H2A-mCherry)*^vo80Tg^, *Tg(NBT:dsRed)*^zf148Tg^. The following transgenic line was established in this study from construct previously reported ^48^: *Tg(zcUAS:PSD95.FingR-TdT-CCR5TC-KRAB(A))* when crossed to a *Tg(chata:gal4)*^mpn202Tg^ was abbreviated as *Chat-PSD95^FingR^*in the manuscript.

### Cloning

The *Tg(zcUAS:PSD95.FingR-TdT-CCR5TC-KRAB(A))*, construct generated in this study was modified from a pTol2-zcUAS:PSD95.FingR-eGFP-CCR5TC-KRAB(A) construct kindly provided by Dr. Joshua Bonkowsky^48^. We replaced eGFP with TdTomato (TdT) in the PSD95.FingR construct as described below. The pTol2-zcUAS:PSD95.FingR-TdT-CCR5TC-KRAB(A) vector was digested by the BglII restriction enzyme. The CCR5TC-KRAB(A) sequence was amplified by PCR from the original vector (forward primer: 5’-gctgtacaagtctaggggagctggcgct-3’, reverse primer: 5’-cccgtggccgtatcttcgcagatctgatctagaggatcataatcagccatac-3’) and TdT sequence was amplified from p3E-TdT vector (forward primer: 5’-taagggtagcggctccagtagatctggggtgagcaagg-3’, reverse primer: 5’-ctcccctagacttgtacagctcgtccatg-3’). The BglII-digested vector and two DNA fragments were integrated by NEBuilder Hifi DNA assembly mix (NEB) to generate the pTol2-zcUAS:PSD95.FingR-TdT-CCR5TC-KRAB(A).

The pDESTTol2CR2-mpeg:mmp14b-HA construct (abbreviated mpeg:mmp14b-HA) was generated in this study by Gateway LR cloning. To prepare the pME-mmp14b gateway donor construct, *mmp14b* gene was amplified by PCR from zebrafish brain cDNA (forward primer: 5’-aaaaagcaggcttagccaccatgatctggagcgggttc-3’, reverse primer: 5’-ttgtacaagaaagctgggttaaccttgtccagtagggag-3’) and the pME vector sequence was amplified from other pME construct (forward primer: 5’-aacccagctttcttgtacaaag-3’, reverse primer: 5’-ggtggctaagcctgcttttttg-3’). The PCR amplified *mmp14b* and pME vector sequences were assembled by NEBuilder Hifi DNA assembly mix to generate the pME-mmp14b. To prepare the p3E-4xHA gateway donor construct, the 4xHA sequence was amplified by PCR from pUB-smHA-KDM5B-MS2 (Addgene, #81085) (forward primer: 5’-ctttcttgtacaaagtggtaggtggatatccctacgacgtg-3’, reverse primer: 5’-tttgtataataaagttgtttaagcgtagtcgggcacgtc-3’) and the p3E vector sequence was amplified from other p3E construct (forward primer: 5’-taaacaactttattatacaaagttgg-3’, reverse primer: 5’-taccactttgtacaagaaag-3’). The PCR amplified 4xHA and p3E vector sequences were assembled by NEBuilder Hifi DNA assembly mix to generate the p3E-4xHA. To generate the pDESTTol2CR2-mpeg:mmp14b-HA vector, the pDESTTol2CR2, p5E-mpeg1.1 (Addgene, #75023), pME-mmp14b, and p3E-4xHA vectors were recombined by gateway LR Clonase enzyme mix (Thermo Fisher Scientific).

To generate the rescue human MMP14-eGFP construct, firstly, silent mutations on MMP14 were introduced which made it resistant to shMMP14_1 (human MMP14_shRNA, Millipore Sigma, TRCN0000429314). We performed PCR (forward primer: 5’-tgCggAgtAccTgacaagttCggggctgagatcaaggccaatgttc-3’, reverse primer: 5’-GaacttgtcAggTacTccGcatcgggggcgcctcatggccttcatg-3’) of the original plasmid pLPCX2-MT1-MMP-eGFP (Addgene, #89819). Then the rescue MMP14 -eGFP sequence containing the silent mutations was amplified by PCR (forward primer: 5’-aggaatttcgacatttggatccatgtctcccgccccaagac-3’, reverse primer: 5’-ataagtttttgttcacgcgtttacttgtacagctcgtccatgc-3’) and integrated to the BamH1-digested vector pLV-EF1A-BSD (kept in house) by Gibson Assembly Cloning Kit (NEB) to generate the rescue MMP14-eGFP construct. The rescue MMP14-eGFP construct was then confirmed by whole plasmid sequencing.

### Embryo microinjection

The injection mix contained 1% Danieau buffer (diluted from 30% Danieau buffer stock: 58 mM NaCl, 0.7 mM KCl, 0.4 mM MgSO_4_, 0.6 mM Ca(NO_3_)_2_, and 5 mM HEPES, adjusted pH to 7.2), 0.1% phenol red, 5 ng/µl TPase mRNA, and 15 ng/µl DNA construct in nuclease-free water as the final concentration. We injected 1 nl of the injection mix into one-cell stage zebrafish embryos with a microinjector. The injected F0 embryos were incubated at 28.5 °C. To generate Tg fish, we injected the pTol2-zcUAS:PSD95.FingR-TdT-CCR5TC-KRAB(A) construct into the *Tg(chata:gal4)* fish. For F0 fish analysis, we injected pDESTTol2CR2-mpeg:mmp14b-HA construct into the *Tg(mpeg:GFP-CAAX)* or *Tg(chat:gal4);Tg(zcUAS:PSD95.FingR-TdT-CCR5TC-KRAB(A))* embryos. The embryos were screened by the red heart marker and used for analysis at 14 dpf.

### CRISPR gene knock-out (KO)

*mmp14b*^-/-^ fish were kindly provided by Dr. Brian Black and generated as described previously^57^. To establish the *bcan*^-/-^ fish, we used the Alt-RTM CRISPR-Cas9 system from IDT. Briefly, gRNA complex was prepared by mixing 100 µM tracrRNA and 100 µM crRNA targeting exon 3 (5’-gcgtacggggaagactcacc-3’) or exon 14 (5’-gacctggtaacatggcgcac-3’) of the *bcan* gene and incubated at 95 °C for 5 min. The gRNAs targeting the *bcan* gene were designed by IDT CRISPR-Cas9 guide RNA design tool and CHOPCHOP. The two gRNA complexes were mixed at 1:1 ratio and mixed with 0.5 µg/µl Cas9 protein diluted in Cas9 working buffer (20 mM HEPES, 150 mM KCl, pH 7.5). The mixture of gRNA complex and Cas9 protein was incubated at 37 °C for 10 minutes to prepare RNP complex. 2 nl of the RNP complex was injected into one-cell stage embryos. Gene deletion was analyzed by PCR (for KO genome, forward primer: 5’-caaagctttaaacactgctgatg-3’, reverse primer: 5’-tcaggtgtttttgttgttgagtc-3’; for WT genome, forward primer: 5’-tgcagttcaatgggttgtgt-3’, reverse primer: 5’-tcaggtgtttttgttgttgagtc-3’). In the F1 generation, we identified heterozygous mutants by fin genotyping and in-crossed the F1 fish. In F2 generation we identified homozygous mutant and wild type fish on this shared genetic background and used age matched larvae for subsequent experiments.

### Immunohistochemistry (IHC) on zebrafish fixed brain sections

Zebrafish at 7, 14, 21, 28, 60, and 90 dpf were fixed overnight at 4 °C in 0.1 M phosphate-buffered 4% paraformaldehyde, cryoprotected with 20% sucrose, and embedded in optimal cutting temperature (OCT) medium (Sakura Finetek USA). IHC was performed on 18-to 25-μm-thick coronal sections collected on a cryostat and mounted onto VWR Superfrost Plus microscope slides (VWR). Sections were washed in phosphate-buffered saline with 0.04% Triton-X (PBST) and incubated with 20% heat-inactivated normal goat serum (NGS; Sigma-Aldrich) in PBST for 1 hour. Primary antibodies were diluted in 3% NGS PBST and applied overnight at 4 °C. Sections were then washed with PBST three times and incubated in secondary antibodies diluted in 3% NGS PBST for 1-2 hours at room temperature. Sections were washed with PBST three times and mounted with DAPI Fluoromount-G (SouthernBiotech). Stained and mounted sections were imaged on an LSM700 confocal microscope (Zeiss) using 20x (NA 0.8) and 63x (NA 1.4) objectives. Images were analyzed with Image J (version 2.14.0).

For HA staining in the mpeg:mmp14b-HA injected fish, signal amplification was performed by using TSA Plus Cyanine 3 system (AKOYA Biosciences). After incubation in the primary antibody, sections were incubated in 3% H_2_O_2_ solution for 30 minutes at room temperature. Sections were washed and secondary antibodies conjugated with HRP were diluted in 3% NGS PBST, applied to sections, and incubated for 1-2 hours at room temperature. After washing sections, Cy3 dye solution (1:50) was applied to sections and incubated for no more than 8 minutes at room temperature. Sections were washed and mounted as described above.

The following primary antibodies were used: anti-GFP chicken polyclonal antibody (Aves Labs GFP-1020, 1:1000), anti-brevican mouse monoclonal antibody (1:100)^88^, Living Colors DsRed polyclonal antibody (Clontech 632496, 1:1000), anti-SV2 mouse monoclonal antibody (DSHB, 1:500), anti-4C4 mouse monoclonal antibody (Gift from Hitchcock lab, 1:200), anti-HA rabbit monoclonal antibody (Cell Signaling Technology 3724T, 1:500). To obtain anti-brevican antibody, SI10-brevican (Addgene, #46300) was transfected into HEK293 cells by Lipofectamine 3000 Transfection reagent (Thermo Scientific) and collected culture media at 48 or 72 hours after transfection. The culture media was concentrated by Amicon Ultra centrifuge filter (Millipore) to 1/10 volume and used for staining at 1:100. The following antibodies were used as secondary antibody at 1:500 dilution: goat anti-chicken Alexa Fluor 488, goat anti-mouse Alexa Fluor 488, goat anti-mouse Alexa Fluor 555, goat anti-mouse Alexa Fluor 647, goat anti-rabbit Alexa Fluor 555 (Invitrogen), goat anti-rabbit HRP (Cell Signaling Technology).

To measure the intensity of brevican staining, the region was selected by the selection tool and measured the mean intensity. Whole hindbrain measurements included an entire coronal section. To measure the ECM intensity from the synaptic region, 100 µm square ROI was placed at the dorsal part of the synaptic region and the mean intensity within ROI was measured. We measured three different regions from 1 section and averaged. We averaged values from two or three sections per fish. Means per fish were plotted and used for statistical comparisons. Values were normalized with the average of control for each experiment.

To measure the intensity of SV2 from the synaptic region, we threshold SV2 signals by thresholding tool (with Li algorithm, Image J). The thresholded region was selected and mean intensity was measured. We measured the intensity from 2-3 sections per fish and averaged. Average per fish was plotted on the graphs and used for statistical comparisons. Values were normalized with the average of control for each experiment.

To count the Chat-PSD95^FingR^ puncta, we randomly took images of single dendrites connected with its cell body. We then cropped the dendrite region and set standard thresholding parameters equivalent across all samples to optimize detection of synaptic puncta. After thresholding, we performed the particle analysis and counted the number of the puncta. We analyzed more than one cell per each fish and generated an average value for each fish when two or more were analyzed. Mean values from each fish were plotted and used for statistical comparisons. Key results were verified with fully blinded quantifications.

Numbers of microglia were counted manually. Values used for plots and statistical analyses based on average microglial numbers per fish from 2-3 sections per fish.

To measure total process length of microglia, z-stack images (0.5 µm step size, 63x objective) encompassing the entire microglia were analyzed using Imaris software (Bitplane, version 9.8.2). The measurement point function was used to calculate the total length of all processes per cell in 3D view. Total process length of each microglia was calculated and plotted.

### Fluorescent in situ hybridization on zebrafish brain sections

Fluorescent in situ hybridization (FISH) experiments were performed using the RNAscope technology (Advanced Cell Diagnostics, ACD) following the manufacturer’s protocol for fixed-frozen tissue, but without the 60 °C incubation and post-fixation steps prior to tissue dehydration. Reagents for RNAscope assay were purchased from ACD. For fish sample preparation, fish were fixed at 14 dpf and OCT blocks were prepared as described in the immunohistochemistry section. We used 14-µm-thick sections collected on cryostat and mounted on VWR Superfrost Plus microscope slides. In this study, we used a C1 probe for zebrafish *mmp14b* (ACD, 1061661-C1). IHC following RNAscope was immediately performed after the last wash of the RNAscope protocol. Sections were incubated in PBS containing 20% NGS at room temperature for 1hr. Primary antibodies were diluted in 3% NGS PBS and incubated at 4 °C overnight. Sections were then washed with PBS three times and secondary antibodies diluted in 3% NGS PBS were applied and incubated at room temperature for 2 hours. Sections were washed with PBS three times and mounted with DAPI Fluoromount-G. Anti-GFP chicken polyclonal antibody and Living Colors DsRed polyclonal antibody were used as primary antibodies at the same concentration as IHC. Anti-chicken Alexa Fluor 488 and anti-rabbit Alexa Fluor 555 were used as secondary antibodies at the same concentration as IHC. Stained sections were imaged on an LSM700 confocal microscope using 63x objective lens. Images were analyzed with Image J.

To count the number of *mmp14b* positive microglia, we used the cell counter function and manually counted the number of mpeg-GFP microglia colocalized with mmp14b puncta. We calculated the percentage of mmp14b+ microglia in total mpeg-GFP+ microglia and averaged 2-3 sections per fish and plotted values for each fish on the graph.

### Brain injections

For the hyaluronidase injection, zebrafish larvae at 10-14 dpf and adult fish at 60 dpf were anesthetized with 0.2 mg/ml tricaine diluted in fish water. For larval injection, larvae were injected with 2 nl of 10 mg/ml hyaluronidase (Invitrogen) or PBS as a vehicle control into the hindbrain ventricle by using a microinjector. The injection was confirmed by seeing a slight expansion of the ventricle. The injected larvae were transferred to fresh fish water after injection for recovery. Larvae were fixed for IHC or used for live imaging at 6 hours post injection (dpi). For adult injection, adult fish were placed in a mold made from 2% agarose gel. A small hole was made in the skull above the hindbrain using a 31 gauge insulin syringe (BD, 328438) and microinjection capillary was inserted from the hole and 40 nl of hyaluronidase or PBS was injected into the hindbrain ventricle. The injected adult fish were transferred to fresh fish water after injection for recovery. Fish were fixed at 6 hpi for IHC. We carefully monitored the fish after injection to confirm that they recovered from anesthesia and did not exhibit abnormal swimming behaviors. Only fish that look healthy were used for further experiments.

### Time lapse live imaging of synapse dynamics (24-72 hrs time lapse)

For the synapse time lapse assay, *Tg(chata:gal4);Tg(zcUAS:PSD95.FingR-TdT-CCR5TC-KRAB(A))* (*Chat-PSD95^FingR^*) fish at 5-14 dpf were used. Larvae were anesthetized with 0.2 mg/ml of tricaine diluted in fish water and mounted in 1.5% low-melting agarose gel on a glass bottom 35-mm dish (MatTek) and covered with fish water containing 0.2 mg/ml tricaine. The still z-stack images were taken on the Nikon or Leica CSU-W1 spinning disk confocal microscope at UCSF Center for Advanced Light Microscopy. We took 40-60 µm z-stack images with 0.5 µm step size. After taking the first image, larvae were carefully removed from agarose gel, placed in a petri dish with fresh fish water, and kept in the fish facility at 28.5 °C with feeding overnight. At the second time point, the larvae were re-mounted in agarose and imaging was repeated as previously described. For the longer time course experiments the same procedure was repeated for up to 72 hours.

The images were processed by ImageJ software as follows: For analysis of average synapse turnover over two time points, two images of the same dendrite were obtained (24 hours apart for most studies, 6 hours apart for pre- and post-hyaluronidase injection experiments, and 30 hours apart for pre- and post-forced swim experiments). We defined Chat-PSD95^FingR^ puncta observed only on the first image as “lost synapses”, puncta observed only on the second image as “new synapses” and puncta visible in both images as “stable synapses”. We carefully compared dendrite shapes and synapses between the first and second images and manually counted the number of the new, lost and stable synapses from the same region of each dendrite, only analyzing regions where Chat-PSD95^FingR^ puncta were clearly defined. We normalized the number of new, lost and stable synapses with the length of dendrites analyzed. Dendrite length was measured from both first and second images and an averaged value was used for the normalization given small differences in fish size and mounting angle between days. At least one cell per fish was analyzed and average values for each fish were used for graphing and statistical analyses.

For merged representative images, we performed image registration by using the bUnwarpJ tool in Image J.

In some cases, the same dendritic segment was imaged repeatedly over 72 hours (t=0 to t=72) to calculate the survival rate of individual synapses. In this case, we defined all synapses present at t=0 as “stable” synapses and synapses newly observed at t=6 as “new” synapses. These two synapse sets were tracked over time to calculate their survival rate. For the representative reconstructed synapse images, we used Imaris software (Bitplane, version 9.8.2). We performed 3D reconstructions using the surfacing function on Imaris. Reconstructed synapses were overlaid with TdT background signals in the dendrites.

To quantify synapse size, we measured the maximum diameter of stable and new synapses at t=6 using the line function in ImageJ. New synapse diameters were normalized to the average of stable synapse diameters for each cell. To quantify the synapse distance to the nearest synapse, we measured the dendrite length from nearest synapses (any synapses including stable and new) by using segmented line function at t=6. The values were normalized by the average of stable synapses for each cell. The values from each synapse were plotted and used for statistical analysis.

### Imaging of microglia synapse contact

*Tg(mpeg:GFP-CAAX);Tg(chat:gal4);Tg(zcUAS:PSD95.FingR-TdT-CCR5TC-KRAB(A))* at 10-12 dpf were mounted for live imaging as described in the previous section. For the hyaluronidase injection experiment, we injected hyaluronidase or PBS as described in the brain injection section and started sample preparation at 4 hpi. Time-lapse imaging was performed on a Nikon CSU-W1 spinning disk/high speed widefield confocal microscope at UCSF Center for Advanced Light Microscopy. We took 40-60 µm z-stack images (step size: 0.5 µm) with 5 minutes intervals for 30 minutes-1 hour. The images were processed by ImageJ software. To count the microglia-synapse contacts, we carefully analyzed single z-stack images and counted the number of Chat-PSD95^FingR^ puncta colocalized with microglial processes. TdT puncta completely merged with GFP signals from microglial processes were defined as “contacted synapses”. To analyze total process movement of microglia, we adapted a previously published method ^89^. Briefly, a single process was chosen per microglia and process length was measured at each time point to calculate the total amount of process tip movement over the entire imaging session to obtain the sum of the total process motility.

### Microglia ablation

We bred fish harboring *Tg(mpeg:gal4);Tg(UAS:NTR-mCherry)* to express nitroreductase (NTR) under control of the *mpeg* promoter. Fresh 5 mM metronidazole (Mtz; Sigma-Aldrich) was prepared in fish water containing 0.2% DMSO before every experiment. Mtz powder was completely dissolved by vigorous shaking. Larvae at 13 dpf were treated with 5 mM Mtz in a petri dish for 24 hours in the dark and fixed at 14 dpf. Control larvae were treated with embryo medium with 0.2% DMSO. For non-transgenic control, we used non-transgenic siblings with no mCherry fluorescence in myeloid cells.

### Light-startle behavior assay

To perform light-startle behavior assay and track fish movement, we used the DanioVision system and EthoVision software (Noldus). Zebrafish larvae at 14 dpf were transferred into a 96-well plate (Cytiva, 7701-1651) and placed in the DanioVision chamber. Fish were habituated in the dark for 30 min before the behavior paradigm started. After habituation, fish movement was tracked according to the following setup: 10 minutes recording before stimuli; 2 second white light stimulus and 2 second dark repeated for 10 times; 10 minutes recording after stimuli. We repeated the same trial for 5 times with no inter-trial intervals. At the conclusion of the experiment all wells were checked for fish viability (normal swimming). Any wells with dead fish were excluded from the analysis.

Startle behavior was analyzed using EthoVision software. To analyze baseline locomotor activity, total distance moved and maximum velocity during the first 10 minutes of recording before the stimuli in trial 1 were calculated from each fish and plotted. To analyze light startle behavior, a plot of velocity over time was binned into 2 second time intervals beginning at 10 seconds before the first stimulus and ending at the end of stimulus sessions. The maximum velocity within each bin was plotted over time. The “maximum response” was defined as the highest velocity achieved over the entire stimulation period.

### Forced swim paradigm

The forced swim paradigm was a modification of previous studies ^68,90^. Zebrafish larvae (n=5-15 fish per dish) at 13 dpf were placed in 60-mm petri dishes with or without a stirrer bar (fisherbrand, 14-513-65) on a stir plate set at 650 r.p.m. for 8 hours per day. The experiment was performed for 2 consecutive days. Larvae were immediately fixed for IHC after the end of the forced swim paradigm at 14 dpf. For synapse turnover assays, images before forced swim were taken in the morning of day 1 and then fish were placed in a petri dish with a stirrer bar for at least 5 hours. On day 2, fish were placed in the petri dish with a stirrer bar for at least 5 hours and then images after forced swim were taken. Experiments and analysis were performed as described in the synapse time lapse assay section above.

### RT-qPCR

For RNA extraction from whole zebrafish larvae, we collected 20 larvae in a tube and homogenized them by syringe and needle. RNA extraction was performed by using QIAGEN RNeasy Mini Kit following manufacturer’s instructions. cDNA synthesis was performed with High Capacity cDNA Reverse Transcription kit (Applied Biosystems). Quantitative PCR reaction was performed in 10 µL of reaction mixture containing cDNA solution, 5 µM forward and reverse primers, fast SYBR Green Master Mix (Applied Biosystems), and nuclease-free water. qPCR was performed by QuantStudio 6 Real-Time PCR system (Applied Biosystems). The primers used for RT-qPCR are *bcan* (forward primer: 5’-gatcgctccgtcagataccc-3’, reverse primer: 5’-gttctccatcgtgccgtagtt-3’) and internal control *ef1a* (forward primer: 5’-ctggaggccagctcaaacat-3’, reverse primer: 5’-atcaagaagagtagtaccgctagcattac-3’). For analysis, we used the ddCt method and showed relative expression levels to each control sample. We collected 20 larvae from three independent fish pairs and each dot in the graph shows each batch of larvae.

For RNA extraction from human iPSC derived microglia, around 1 million cells were collected per group in a 1.5 mL tube and RNA extraction was performed using QIAGEN RNeasy Plus Mini Kit following manufacturer’s instructions. cDNA synthesis was then performed with Primescript RT Reagent Kit (Takara Bio) or Superscript IV First Strand Synthesis System (Invitrogen). Quantitative PCR reaction was performed in 10 µL of reaction mixture containing cDNA solution, 1 µM forward and reverse primers, fast SYBR Green Master Mix (Applied Biosystems), and nuclease-free water. qPCR was performed by QuantStudio 6 Real-Time PCR System (Applied Biosystems). The primers used for RT-qPCR are MMP14 (forward primer: 5’-gcagaagttttacggcttgcaa-3’, reverse primer: 5’-ccttcgaacattggccttgat-3’) and internal control GAPDH (forward primer: 5’-tcaacgaccccttcattgac-3’, reverse primer: 5’-atgcagggatgatgttctgg-3’). Relative expression levels were analyzed to each control sample. We collected cells from three independent experiments and each dot in the graph shows the mean value from an independent experiment.

### Bulk RNA-sequencing analysis of metalloprotease genes

The bulk RNA-sequencing data of zebrafish microglia were from our previous study^40^. Briefly, 28 dpf zebrafish brains were microdissected into optic tectum, midbrain, and hindbrain, and microglia were isolated by flow cytometry using fluorescence of *mpeg1.1:EGFP*, followed by library generation and sequencing. The data was screened for all genes beginning with “*mmp*” “*adam*” and “*adamts*.” Genes with at least 10 counts in >1 sample were defined as detectable and plotted on a heatmap of raw count values.

### Human induced pluripotent stem cells induced tri-culture

Human induced pluripotent stem cells (iPSCs, male donor, WTC11 background) were engineered and differentiated, as previously described, with a doxycycline-inducible NGN2 cassette for iNeuron differentiation^91^, doxycycline-inducible SOX9/NFIA cassette for iAstrocyte differentiation^61^, or doxycycline-inducible six transcription factor cassette (6TF: MAFB, IRF5, IRF8, PU.1, CEBPα, CEBPβ) for iTF-Microglia differentiation^62^ in Martin Kampmann lab at UCSF. Briefly, for the tri-culture system in this study, iPSC-derived astrocytes (at 20 days treatment with ScienCell Astrocyte Medium (ScienCell, 1801) were thawed and seeded at 30,000 cells/cm^2^ on Matrigel (Corning, 356231)-coated plates. For the following description, day counting will be based on the seeding of iAstrocytes. On the next day (Day 1 for triculture generation), pre-differentiated iNeurons were seeded on top of iAstrocytes at a density of 60,000 cells/cm^2^ and differentiated for 14 days in the following medium: BrainPhys (STEMCELL Technologies, 05791) as the base, 0.5× B27 Supplement Minus Vitamin A (Life Technologies, 12587010), 1× N2 Supplement (Life Technologies, 17502-048), 10 ng/mL NT-3 (PeproTech, 450-03), 10 ng/mL BDNF (PeproTech, 450-02), 1 μg/mL mouse laminin (Thermo Fisher Scientific, 23017-015), and 2 μg/mL doxycycline (Takara Bio, 631311). The medium was fully replaced at Day 4 with doxycycline and half-replaced without doxycycline every 2-3 days since Day 7. At Day 7, iPSC-6TF was differentiated into microglia in parallel as the following: iPSC-6TF was seeded onto Poly-d-Lysine + Matrigel double-coated plates in the following medium: Essential 8 Basal Medium (Gibco, A15169-01) as a base, 10 nM ROCK inhibitor (Fisher Scientific, 125410) and 2 μg/mL doxycycline. Two days later at Day 9, medium was replaced with the microglia differentiation medium as following: Advanced DMEM/F12 Medium (Gibco, 12634-010) as a base, 1 × Antibiotic-Antimycotic (Gibco, 15240-062), 1 × GlutaMAX (Gibco, 35050-061), 2 μg/mL doxycycline, 100 ng/mL Human IL-34 (Peprotech, 200-34) and 10 ng/mL Human GM-CSF (Peprotech, 300-03). Another two days later at Day 11, medium was replaced to the microglia differentiation medium with addition of 50 ng/mL Human M-CSF (Peprotech, 300-25) and 50 ng/ml Human TGFB1 (Peprotech, 100-21C). At Day15, TrypLE (Gibco, 12605010) was used to dissociate the microglia from the plate. Then dissociated microglia were added at 30,000 cells/cm^2^ to the astrocyte and neuron co-culture to form the triculture system. At Day 16, the triculture system was then used in western blot and immunofluorescence as described later.

### Process length analysis through immunofluorescence

Cells were washed carefully with PBS once and then fixed with 4% paraformaldehyde for 30 minutes at room temperature. After PBS washing for three times, cells were permeabilized with SuperBlock buffer (Thermo Scientific, 37515) containing 0.1% Triton X-100 for 1 hour at room temperature. Cells were then incubated with primary antibodies diluted in PBS at 4 °C overnight. Primary antibodies used in this study were as follows: chicken anti-MAP2 polyclonal antibody (1:500, Invitrogen, PA116751), rabbit anti-IBA1 (1:200, Wako Chemicals 019-19741), mouse anti-S100β antibody (1:200, Sigma-Aldrich, S2532). On the next day, after PBST washing for three times, cells were incubated with secondary antibodies diluted in PBS for 2 hrs at room temperature. After that, cells were washed with PBST three times and washed with ddH2O once. Cells were then mounted with DAPI Fluoromount-G Mounting Medium (SouthernBiotech, 0100-20) and images were taken using a Leica Widefield microscope. For microglia (IBA1+) with processes, their process length was measured using ImageJ.

### Protein detection by western blotting

For Brevican detection, cell culture medium was collected in 1.5 mL Ep tube separately and centrifuged at 13,000 rpm for 1 minute at 4°C to get rid of potential debris. Then the supernatant was concentrated tenfold in the Amicon Centrifugal Filter Unit (10kD, EMD Millipore, UFC501096). 4x Laemmli Sample Buffer (Bio-Rad, 1610747) was added to concentrated supernatant and boiled at 100°C for 10 minutes. At the same time, cells were collected in 1x Laemmli Sample Buffer separately and boiled at 100°C for 10 minutes. For western blotting, 20 μg of total proteins were loaded into 4–15% Mini-PROTEAN TGX Precast Protein Gels (Bio-Rad, 4561086). Subsequently, the gels were transferred into Immun-Blot PVDF Membrane (Bio-Rad, 1620255). The membranes were blocked by 5% milk (Bio-Rad, 1706404), followed by incubation with primary antibodies at 4°C overnight. Primary antibodies used were as follows: rabbit anti-Brevican polyclonal antibody (1:1000, Thermo Fisher Scientific, PA552477), and mouse anti-MAP2 monoclonal antibody (1:1000, Thermo Scientific, 13-1500) as inner control. On the next day, the membranes were washed three times with TBST (Tris-buffered saline with 0.1% Tween detergent, Bio-Rad, 1706435) and then incubated with secondary antibodies for 2 hrs at room temperature. The membranes were then washed three times with TBST and developed in Clarity Western ECL Substrate (Bio-Rad, 1705060). Images were taken using ChemiDoc Imaging Systems and analyzed using ImageJ. Four independent experiments were performed and each dot in the graph represents the relative intensity from each independent experiment. For MMP14 detection, microglia cells were collected in 1x Laemmli Sample Buffer and boiled at 100°C for 10 minutes. The same procedure outlined above was followed, except that the primary antibodies used were rabbit anti-MMP14 monoclonal antibody (1:1000, Cell Signaling Technology, 26424S) and rabbit anti-β-Actin monoclonal antibody (1:1000, Cell Signaling Technology, 4970S) as inner control.

### Gene expression analysis of human fetal microglia and iPSC-derived microglia

Bulk RNAseq data of human fetal microglia and iPSC-derived microglia were obtained from the previous study^59^. Within each group, genes are ranked by their mean TPM across samples. Scatter plots showing the top 8000 of gene expression rank were generated using matplotlib and python.

### Sample Preparation for Proteomics

Protein samples were denatured by addition of 50 µl of 6M guanidine hydrochloride,100mM Tris pH 8. Proteins were then reduced and alkylated by addition of Tris(2-carboxyethyl)phosphine (TCEP) and 2-chloroacetamide (CAA) to a final concentration of 10 mM and 40 mM, respectively, before incubation at 95 °C with 700 rpm for 7 minutes. Guanidine was diluted 1:5 by the addition of 200 µl of 100mM Tris pH 8. Proteins were digested by adding 1 µg of trypsin to each sample followed by incubation overnight at 37 °C with 800 rpm shaking.

Digested peptides were acidified by the addition of 50 µl of 10% trifluoroacetic acid (TFA) before desalting using NEST UltraMicrospin tips attached to a vacuum manifold. Each tip was activated with 100 µl 80% acetonitrile (ACN), 0.1% TFA, then equilibrated with three washes of 150 µl of 0.1% TFA. Peptides were loaded onto the tips, before washing four times with 150 µl of 0.1% TFA. Peptides were eluted by centrifugation after addition of 150 µl 50% ACN, 0.25% formic acid (FA). Eluted peptides were dried down in Speed-Vac concentrator before being resuspended in 60 µl 0.1% FA. Resuspended peptide samples were filtered by centrifugation (Millipore 0.45 µm hydrophilic low protein binding Durapore Membrane) and diluted 1:10 before 3 µl was injected onto the LC-MS system.

### Liquid Chromatography Mass Spectrometry

Samples were analyzed on an Orbitrap Exploris 480 mass spectrometry system (Thermo Fisher Scientific) equipped with Neo Vanquish ultra-high pressure liquid chromatography system (Thermo Fisher Scientific) interfaced via a Nanospray Flex source. Separation was performed using a 15 cm long PepSep column with a 150 µm inner diameter packed with 1.5 µm Reprosil C18 particles. Mobile phase A consisted of 0.1% FA, and mobile phase B consisted of 0.1% FA, 80% ACN. Peptide samples were separated by a gradient of 3% to 28% mobile phase B over 67 minutes followed by an increase to 40% B over 5 minutes. Chromatographic gradients ended with a wash at 95% B for 8 minutes. The flow rate was 600 nL/minute throughout, outside of sample loading, which was pressure controlled, with a max pressure of 300 bar. Spectra were acquired in a data-independent manner (DIA) with a full scan across 350-1050 m/z in the Orbitrap at 60,000 resolving power (at 200 m/z) with a normalized AGC target of 300%, an RF lens setting of 40%, and a maximum ion injection time of “Auto”. DIA isolation windows were generated automatically, covering 350-1050 m/z space with 20 m/z wide windows and 0.5 m/z overlap. Peptide ions were fragmented using a normalized HCD collision energy of 28%, with an AGC target of “standard”, a maximum injection time of 40 ms, with scan range mode set to “auto”, and a resolving power of 15,000. Gas phase fraction (GPF) data was collected from a pooled sample as described by Pino et al^92^ to supplement our existing spectral libraries. Resolution, fragmentation and chromatographic parameters were identical to DIA runs with full scans covering 110 m/z windows with 5 m/z overlap (e.g. 345-455, 445-555, etc.) and fragment scan windows of 4 m/z.

### Mass Spectrometry Data Searching and Analysis

DIA data files were searched in Spectronaut (version 19.5) against a spectral library developed in-house from the full, reviewed human proteome with isoforms (downloaded from UniprotKB May 30, 2023) and supplemented with our GPF acquisition files. Search was performed with default settings, except cross run normalization, which was not used. Results were output in the MSStats Format. Peptide features were summarized into protein abundances and the two groups were compared using the dataProcess and groupComparison functions of the MSstats^93^ package (version 4.12.0), respectively. The group comparison results were used to generate the significance and fold change for proteins depicted in Figure 4. For the proteomics results, see Extended Table 1,2.

### Computational analysis

We fitted models for the synapse population dynamics to the synapse lifetime imaging data, in order to determine if the observations are better described by a single or two (dynamic and stable) internal states of the synapses as well as to estimate the four key model parameters (synapse birth rate, decay rate of dynamic synapses, transition rate from dynamic to stable synapses and the decay rate of stable synapses) for *mmp14^+/+^* and *mmp14b^-/-^* conditions, as shown in Figure 7. See Supplemental File 1 for mathematical derivation of the models and their predictions of experimental observations and numerical fitting.

### Statistical analysis

Graphpad prism 10.3.0 was used for all statistical analysis. Statistical tests used, number of n replicates and p values are described in figures, text and figure legends. All statistical analysis unless otherwise noted were performed on means of multiple technical replicates per animal. For comparisons of two groups, we used t-test with Welch’s correction to correct unequal variance. Data sets with one than two groups were analyzed with one-way or two-way ANOVA.

