## Supplemental File 1 for "Extracellular matrix proteolysis maintains synapse plasticity during brain development"

### Supplement: Modeling of developmental synapse dynamics

Here we describe the birth-decay processes models used to describe the developmental dynamics of synaptic life times observed in the experiments. We provide derivations of the model predictions, including predictions for the experimental measurements which are used to fit the parameter to the data. In our modeling approach, we assume that synapses are created via a random process and then transition with certain probabilities through one or multiple internal states before they disappear. We show that a model with only a single internal state cannot explain the experimental data. However, already a model with two states of synapses, including a newly generated state as well as a stable state in which the survival rate is higher, describes the experimental data well and fits 21 data points with only 4 parameter. Our analysis indicates that synapses come in two flavors: a dynamic pool into which synapses are born and from which they decays faster, and a stable pool to which new synapses can transition and from which decay is much slower. By fitting these two states models to control and knock-out data we find that key parameters change in accordance with the main interpretation of our data as described in the main manuscript.

In the following we will introduce the birth-decay processes model for developmental synaptic dynamics, derive key predictions of the models, including predictions for the experimental measurements, and describe our fitting procedures to the experimental data. As the one state model can be viewed as a special case of the two state model we consider the two state model first and specialize to the one state model in the end. If not stated otherwise we model the number or density of synapses on a fixed segment of a dendrite.

#### Two state synapse model

Our modeling approach is based on birth-decay stochastic process models that have been previously used to describe nuclear decay or dynamics of epidemics [1, 2]. We assume two types of synapses:

- "new" synapses denoted by $n$ of recently generated and
- "stable" synapses $s$ that transitioned to a more stable state.

New synapses are generated via a homogeneous Poisson process with a birth rate $r_{\emptyset,n}$. A single new synapse will transition to stable synapses with a rate $g_{n,s}$ or decay with a rate $g_{n,d}$. Given $N(t)$ new synapses at time $t$, the total transition rates are thus

$$r_{n,s}(t) = g_{n,s}N(t) \qquad r_{n,d}(t) = r_{n,d}N(t)$$

The total loss of new synapses is

$$r_{n,-}(t) = r_{n,d}(t) + r_{n,s}(t) = (g_{n,s} + g_{n,d})\, N(t) = g_{n,-}N(t)$$

were we introduced $g_{n,-} = g_{n,s} + g_{n,d}$, the total loss rate of new synapses. Each stable synapses will decay with a typically smaller rate $g_{s,d}$. Given $S(t)$ stable synapses the decay rate is

$$r_{s,d}(t) = g_{s,d}S(t)$$

The number of new and stable synapses are random variables $N(t)$ and $S(t)$ that evolve according to the above stochastic dynamics and these two numbers fully characterize the state of the system. The probability to be in a specific state at time $t$ is denoted as

$$p(n, s, t) = \mathbb{P}(N(t) = n, S(t) = s)$$

The model is fully specified by its four parameters

$$\theta = \left(r_{\emptyset,n}, g_{n,d}, g_{n,s}, g_{s,d}\right)$$

as well as the distribution of initial conditions $p(n, s, t_0) = p_0(n, s)$. We denote initial conditions centered on a deterministic state as $N(t_0) = n_0$ and $S(t_0) = s_0$.

#### Master equation

Note that the probability for more than one event to happen in the time interval $dt$ is $\mathcal{O}(dt)$. Thus given a state $(n, s)$ at time $t$ we have the following probabilities for single events to happen within $dt$ up to that order:

$$
\begin{aligned}
q_{n,s\to n+1,s} &= r_{\emptyset,n}dt + \mathcal{O}(dt) \\
q_{n,s\to n-1,s} &= ng_{n,d}dt + \mathcal{O}(dt) \\
q_{n,s\to n-1,s+1} &= ng_{n,s}dt + \mathcal{O}(dt) \\
q_{n,s\to n,s-1} &= sg_{s,d}dt + \mathcal{O}(dt)
\end{aligned}
$$

and thus for no event to happen

$$
q_{n,s\to n,s} = 1 - q_{n,s\to n+1,s} - q_{n,s\to n-1,s} - q_{n,s\to n-1,s+1} - q_{n,s\to n,s-1}
$$

Thus, the probabilities $p(n, s, t + dt)$ for the system to be in a state $(n, s)$ at a time $t + dt$ is given by:

$$
\begin{aligned}
p(n,s,t+dt) =& q_{n,s\to n,s}p(n,s,t) + q_{n-1,s\to n,s}p(n-1,s,t) + q_{n+1,s\to n,s}p(n+1,s,t) \\
&+ q_{n+1,s-1\to n,s}p(n+1,s-1,t) + q_{n,s+1\to n,s}p(n,s+1,t) + \mathcal{O}(dt)
\end{aligned}
$$

We can rearrange and and take the limit $dt \to 0$ to obtain the master equations:

$$
\begin{aligned}
\partial_t p(n,s,t) =& -\left(r_{\emptyset,n} + ng_{n,d} + ng_{n,s} + sg_{s,d}\right)p(n,s,t) + r_{\emptyset,n}p(n-1,s,t) + (n+1)g_{n,d}p(n+1,s,t) \\
&+ (n+1)g_{n,s}p(n+1,s-1,t) + (s+1)g_{s,d}p(n,s+1,t)
\end{aligned} \tag{1}
$$

#### Average dynamics

For the average number of new $\bar{n}$ and stable $\bar{s}$ synapses

$$
\begin{aligned}
\bar{n}(t) &= \mathbb{E}(N(t)) = \sum_n np(n,t) \\
\bar{s}(t) &= \mathbb{E}(S(t)) = \sum_s sp(s,t)
\end{aligned} \tag{2}
$$

we can derive their evolution equations from the master equations (1) as :

$$
\begin{aligned}
\partial_t\bar{n}(t) &= r_{\emptyset,n} - \bar{n}(t)g_{n,-} \\
\partial_t\bar{s}(t) &= \bar{n}(t)g_{n,s} - \bar{s}(t)g_{s,d}
\end{aligned}
$$

which for the initial conditions

$$
\bar{n}(t_0) = \bar{n}_0 \qquad \bar{s}(t_0) = \bar{s}_0
$$

can be integrated to yield

$$
\begin{aligned}
\bar{n}(t) &= \bar{n}_\infty + (\bar{n}_0 - \bar{n}_\infty)e^{-g_{n,-}(t-t_0)} \\
\bar{s}(t) &= e^{-g_{s,d}(t-t_0)}\left(\bar{s}_0 + \int_{t_0}^{t} e^{g_{s,d}(s-t_0)}g_{n,s}\bar{n}(s)ds\right)
\end{aligned} \tag{3}
$$

with steady state solution $(t \to \infty)$

$$
\bar{n}_\infty = \frac{r_{\emptyset,n}}{g_{n,-}} \qquad \bar{s}_\infty = \bar{n}_\infty\frac{g_{n,s}}{g_{s,d}} = \frac{r_{\emptyset,n}}{g_{n,-}}\frac{g_{n,s}}{g_{s,d}} \tag{4}
$$

We can solve for $s$ explicitly

$$
\bar{s}(t) = \bar{s}_\infty + (\bar{s}_0 - \bar{s}_\infty)e^{-g_{s,d}(t-t_0)} + (\bar{n}_0 - \bar{n}_\infty)\frac{g_{n,s}}{g_{n,-} - g_{s,d}}\left(e^{-g_{s,d}(t-t_0)} - e^{-g_{n,-}(t-t_0)}\right)
$$

with a direct interpretation: The first term is the steady state, the second term captures initial deviations of stable synapses from the steady state while the last term captures deviations in the new state which first transition to the stable state and then decay form there.

This concludes the derivation of the population and average dynamics for the synapses which we can use to derive expressions for the average observed synapse numbers in experiments.

#### Average observations

##### Experimental Measures

Experimentally we cannot monitor the synapses continuously, and thus only observe them a discrete points in time. We here derive the predictions from our model for the average observed number of synapses.

More concretely, as in experiments, we measure occurrences of synapses at $K \in \mathbb{N}$ observation times $t_k$, $k \in \{1, 2, \ldots, K\}$ with $t_k < t_{k+1}$. For later mathematical convenience we define further observation times $t_0 = -\infty$ and $t_{K+1} = \infty$ to absorb boundary cases and discuss their role below. In our experiments we have $K = 6$ and observation times at $t_1 = 0\text{h}$, $t_2 = 6\text{h}$, $t_3 = 12\text{h}$, $t_4 = 24\text{h}$, $t_5 = 48\text{h}$ and $t_6 = 72\text{h}$.

For each synapse it is determined at which observation time $t_f$ it first appeared (i.e. it was not observed at the previous observation time $t_{f-1}$) as well as at what observation time $t_l$ it was seen last (i.e. not observed at $t_{l+1}$ anymore). Then the number of synapses that were observed from $t_f$ to $t_l$ for all pairs of $f \leq l$ are counted. We denote these numbers of synapses *observed* between $t_f$ and $t_l$ by

$$O_{f,l} = O\left(t_{f-1}, t_f, t_l, t_{l+1}\right)$$

where we emphasized on the right hand side of the equation that $O_{f,l}$ depends on four observation time points. From this, the *total number of observed synapses* at a single observation time $t_k$ denoted by $M_k = M(t_k)$ can be derived as:

$$M_k = \sum_{f=0}^{k} \sum_{l=k}^{K+1} O_{f,l}$$

For the synapses that *appeared* between $t_{f-1}$ and $t_f$ denoted by $A_f = A(t_{f-1}, t_f)$ we have

$$A_f = \sum_{l=f}^{K} O_{f,l}$$

Finally, we define the number of synapses that first appeared at $t_f$ and *remained* observable until at least $t_r \geq t_f$ as $R_{f,r} = R\left(t_{f-1}, t_f, t_r\right)$. We have

$$R_{f,r} = \sum_{l=r}^{K+1} O_{f,l} \leq R_{f,f}$$

Inverting this relationship we obtain

$$O_{f,l} = R_{f,l} - R_{f,l+1} \tag{5}$$

Thus, deriving the measures $R_{f,l}$ from our model will enable us to fit it to our experimental observations $O_{f,l}$.

##### Model predictions

Here we will use the average developmental synapse dynamics (3) to predict the average numbers of observed synapses. As we have no experimental access to the internal state of the synapses at any instance of time $t_k$, the total number of synapses we observe is given by

$$M_k = N(t_k) + S(t_k)$$

and on average

$$\bar{m}_k = \bar{n}(t_k) + \bar{s}(t_k)$$

with $\bar{n}$ and $\bar{s}$ given by (3) which depend on initial conditions. Under the assumption that the system is in a steady state this reduces to

$$\bar{m}_k = \bar{n}_\infty + \bar{s}_\infty = \bar{m}_\infty$$

independent of the observation time $t_k$ and initial synapse counts.

For the number of synapses $A_f$ that appeared for the first time at $t_f$ and thus were not observed at the previous observation time $t_{f-1}$ we can use (3) with $n_0 = s_0 = 0$ and $t_0 = t_{f-1}$, $t = t_f$ to obtain for the average appearance of new and stable synapses

$$\bar{a}_f^{(n)} = \bar{n}_\infty \left(1 - e^{-g_{n,-}(t_f - t_{f-1})}\right) \tag{6}$$
$$\bar{a}_f^{(s)} = \bar{s}_\infty \left(1 - e^{-g_{s,d}(t_f - t_{f-1})}\right) - \bar{n}_\infty \frac{g_{n,s}}{g_{n,-} - g_{s,d}} \left(e^{-g_{s,d}(t_f - t_{f-1})} - e^{-g_{n,-}(t_f - t_{f-1})}\right)$$

and thus in total

$$\bar{a}_f = \bar{a}_f^{(n)} + \bar{a}_f^{(s)}$$

Note that for $f \geq 2$ these observations of appearing synapses do not depend on the initial state and are thus very useful types of observations to calibrate the model.

We can further calculate the change in the number of synapses that appeared at $t_f$ and remain visible in a subsequent observation at time $t_r$. As we focus on the already appeared synapses at $t_f$, we can ignore newly generated synapses after $t_f$. In this case we can set $r_{\emptyset,n} = 0$ in our model to describe the dynamics and use solutions (2) for the average dynamics. We have $\bar{n}_\infty = \bar{s}_\infty = 0$ in this case as all synapses will decay eventually and our initial conditions are given by the average number of synapses appearing at $t_f$, (6), i.e. $n_0 = \bar{a}_f^{(n)}$ and $s_0 = \bar{a}_f^{(s)}$. This results in an average number of remaining synapses in the new and stable states given by

$$\bar{r}_{f,r}^{(n)} = \bar{a}_f^{(n)} e^{-g_{n,-}(t_r - t_f)}$$
$$\bar{r}_{f,r}^{(s)} = \bar{a}_f^{(s)} e^{-g_{s,d}(t_r - t_f)} + \bar{a}_f^{(n)} \frac{g_{n,s}}{g_{n,-} - g_{s,d}} \left(e^{-g_{s,d}(t_r - t_f)} - e^{-g_{n,-}(t_r - t_f)}\right) \tag{7}$$

and for the average total number of remaining synapses

$$\begin{aligned}\bar{r}_{f,r} =& \bar{r}_{f,r}^{(n)} + \bar{r}_{f,r}^{(s)} \\ =& \bar{n}_\infty \left(e^{-g_{n,-}(t_r - t_f)} - e^{-g_{n,-}(t_r - t_{f-1})}\right) + \bar{s}_\infty \left(e^{-g_{s,d}(t_r - t_f)} - e^{-g_{s,d}(t_r - t_{f-1})}\right) \\ &+ \bar{n}_\infty \frac{g_{n,s}}{g_{n,-} - g_{s,d}} \left(e^{-g_{s,d}(t_r - t_f)} - e^{-g_{n,-}(t_r - t_f)}\right)\end{aligned}$$

These average expressions for the observations in the model can now be used to fit the model parameter $\theta$ to the experimental data. In particular, we can connect the results here to the experimental observations $O_{f,l}$ using (5). On average

$$\bar{o}_{f,l} = \bar{r}_{f,l} - \bar{r}_{f,l+1}$$

which are independent of any additional assumptions for $2 \leq f \leq l$. Like the total number of synapses $\bar{m}_k$, the initial observations $\bar{o}_{1,l}$ will depend on additional assumptions about the initial state. Under the steady state assumption we can take the limit $t_0 \to -\infty$ as indicated above.

### One State Model

For the one state model we simply forbid transitions to the stable state and set $g_{n,s} = 0$ and allow no stable types of synapses initially $s_0 = 0$. It follows $\bar{s}_\infty = 0$ and inserting these values in the above equations we obtain the predictions for the measurements in the one state model.

### Model Fitting

To fit our model to the data we performed a non-linear model fit using least squares optimization between the model predictions and the experimental data. To compare wild type and knock-out data, we normalized our data to the total dendrite length in each case. We then used a Levenberg–Marquardt algorithm[3, 4] implemented in python[5], to perform the numerical optimization. We initialized parameters from a manual fit. To obtain estimates for the variability of these parameter estimates we randomly sub-sampled the observation of the synapse life times at 90%, and normalized the resulting counts by 90% of the corresponding measured total dendritic length. We repeated the procedure 100 times and used the resulting fitted parameter values to estimate their mean and standard deviation. We used the same data to perform a Mann Whitney U-test to test if parameters are statistically the same or different between the wild type and knock out and report the p-values of this test.

#### Discussion

Our results indicate that a two state model is sufficient to fit the observed data reasonable well, compared to a one state model. In addition to the results shown here we derived lengthy expressions for the distributions of the experimental observations and performed inference on the parameters with similar results. We also studied models in which the synaptic decay is a exponentially decreasing function of the synapse life time requiring no explicit state transitions. We will present these results in a future study.

Our python code of the proposed model and data fitting routines will be made available on github [6] or can be obtained upon reasonable request.

[6] https://github.com/ChristophKirst
